# Counterions, Polarization and Pore Hydration Relay Drive Ion Permeation through Hydrophobic Pore Formed by the MERS Coronavirus E Protein

**DOI:** 10.64898/2026.09.03.749198

**Authors:** Zhongyi Wan, Mei Hong, Qiang Cui

**Author notes:** Author contributions: Z.W., M.H. and Q.C. designed research; Z.W. performed research; Z.W. and Q.C. analyzed data; and Z.W., M.H. and Q.C. wrote the paper. The authors do no have competing interests.

## Abstract

Hydrophobic pores are generally expected to exclude ions because the large desolvation penalty associated with ion transfer cannot be compensated within a nonpolar environment. The envelope (E) protein of Middle East Respiratory Syndrome coronavirus (MERS-CoV) presents a striking exception, conducting cations with high efficiency despite forming a predominantly hydrophobic pore. To resolve this apparent paradox, we combine finite-temperature string simulations with additive and polarizable force fields to determine the minimum free-energy pathways for ion permeation through the MERS E channel. The simulations reveal a cooperative permeation mechanism in which transient counterion association, localized pore hydration, and electronic polarization jointly reduce the desolvation penalty. Rather than acting as passive spectators, counterions transiently chaperone the permeating cation before dissociating within the channel, thereby lowering the energetic cost of charge desolvation while preserving net ionic conductance. Explicit electronic polarization further reshapes the pore hydration landscape and cation–*π* interactions, leading to substantially reduced free-energy barriers and a permeation mechanism in substantially better agreement with experiment. These results suggest that efficient ion transport through hydrophobic pores arises from the cooperative interplay between electrostatics, polarization and hydration rather than complete pore wetting, providing both a mechanistic framework for MERS E protein function and a general physical principle for ion transport through biological and synthetic hydrophobic nanopores.

## 1 Introduction

Efficient transport through hydrophobic nanopores represents a fundamental challenge in biology, chemistry, and nanotechnology[1–3]. While water molecules readily permeate hydrophobic channels such as carbon nanotubes [4, 5], ions face a fundamentally different thermodynamic barrier because transferring a charged species from bulk water into a low-dielectric environment requires partial or complete desolvation. Consequently, hydrophobicity is generally considered incompatible with efficient ion conduction and is frequently invoked as the basis of hydrophobic gating in biological ion channels [6–10]. Yet a growing number of natural and synthetic nanopores appear capable of transporting ions through predominantly hydrophobic pathways[11], raising the fundamental question of how the associated desolvation penalty is overcome.

Coronavirus envelope (E) proteins exemplify these structural and functional transport paradoxes in viroporin biophysics. The small pentameric membrane proteins function as critical virulence factors that form ion-conducting channels essential for viral pathogenicity, making them highly attractive antiviral targets across the coronavirus family. Electrophysiological studies have demonstrated substantial cation conductance for both SARS-CoV-2 and MERS-CoV E proteins. The MERS E protein conducts potassium ions (K^+^) with a remarkably high conductance of 113 picosiemens (pS) at neutral pH [14]. Yet recent solidstate NMR (ssNMR) structures[12, 13] reveal that the MERS E transmembrane domain (ETM) features a pore (see Figure 1A) lacking the extensive polar lining that forms the cation recruiting/relaying network such as the carbonyls in the selectivity filter of canonical potassium channels[15, 16] and the N-terminal threonine network in the SARS-CoV-2 E protein[17]. Instead, it relies on an extraordinarily hydrophobic environment dominated by phenylalanine clusters (F12, F16, F17, F19, and F33)[17] and a non-canonical conformation-selection mechanism where K^+^ binding merely homogenizes pre-existing states rather than inducing open-state transitions [18]. These observations—combined with mutagenesis showing that substituting the N-terminal N15 or F17 abolishes conductance, whereas the C-terminal F33 likely regulates transport via cation–*π* interactions—suggest that ion permeation proceeds through mechanisms fundamentally different from those established for canonical ion channels.

**Figure 1:**
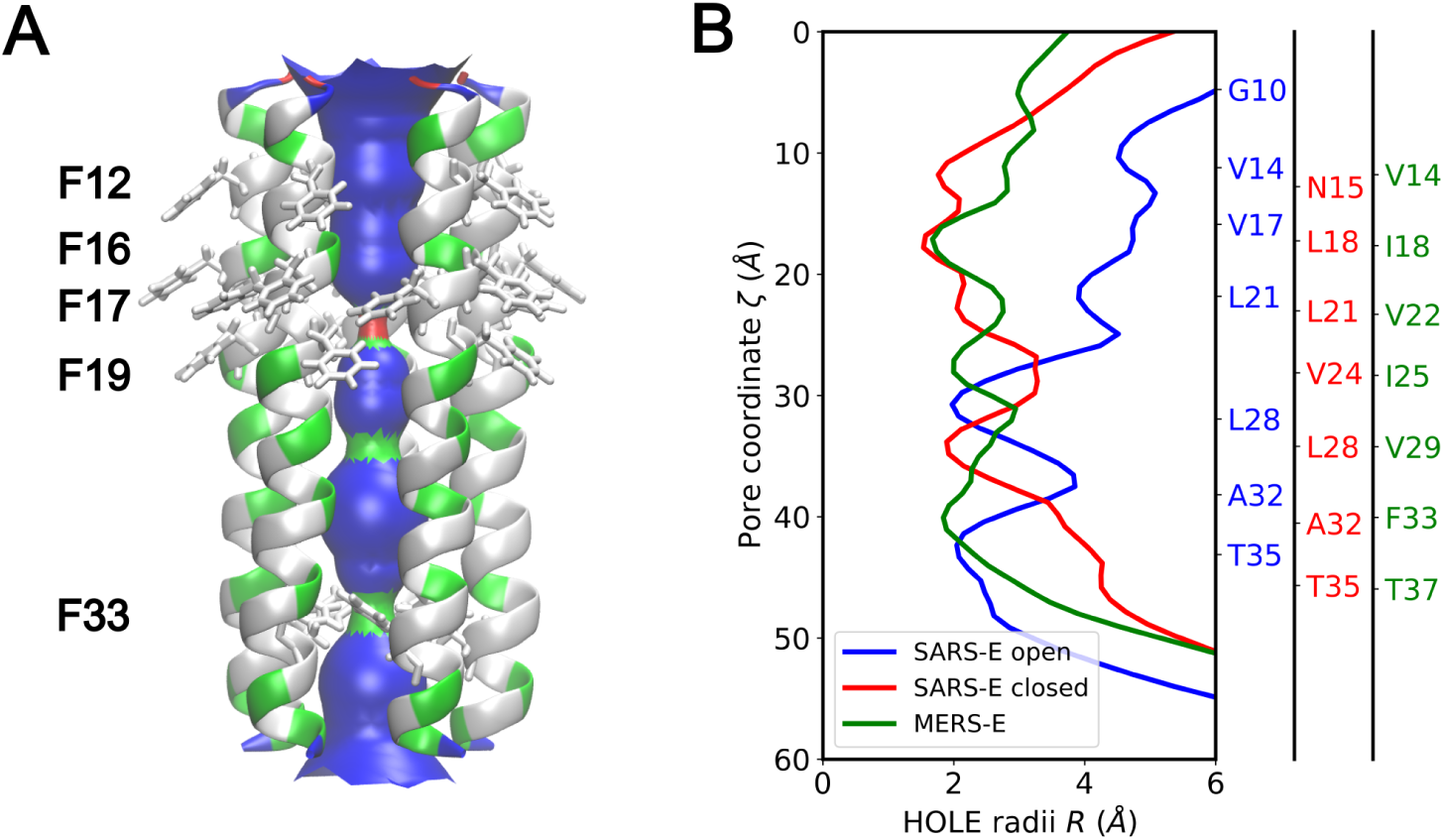
Structural and pore radius analysis of the MERS-CoV E protein. (A) Solid-state NMR (ssNMR) structure of the MERS-CoV envelope transmembrane (ETM) domain shown alongside its calculated hole radii surface. Polar and hydrophobic residues are color-coded in green and gray, respectively, with phenylalanine residues highlighted. (B) HOLE radii profiles of the MERS ETM (PDB ID: 9DCJ) compared against the open (PDB ID: 8SUZ[12]) and closed (PDB ID: 7K3G[13]) states of the SARS-CoV-2 ETM. Corresponding residue C*_α_* positions are indicated along the channel axis.

To resolve how this unconventional hydrophobic architecture supports efficient transport, we employ all-atom molecular dynamics (MD) and free energy simulations. Specifically, finite-temperature string calculations reveal evidence of a counterion-assisted permeation mechanism, explaining how cations traverse the narrow, hydrophobic pore (Figure 1). Analysis of the simulation results using both fixed-charge and polarizable force fields highlights the cooperative roles of electronic polarization, counterions and aromatic/polar residues in modulating the pore hydration landscape to control ion transport. By bridging the gap between static structural data and functional conductance, these simulations providing key insights into this non-canonical transport phenomenon and lay a foundation for E-protein-targeted drug development. The findings also suggest a general physical framework for understanding ion transport through biological and synthetic hydrophobic nanopores.

## 2 Results and Discussion

### 2.1 Characterizing Permeation Pathways via the Finite-Temperature String Method

Since the computational approach we take to analyze the permeation mechanism for the MERS-E protein is somewhat unique for ion channels, we first summarize the motivation and key ingredient of the simulation methodology. Exploratory simulations made it clear that ion permeation in these hydrophobic channels cannot be adequately described by the typical one-dimensional potential of mean force profile along the pore axis since multiple processes in addition to cation displacement are implicated. Accordingly, the finite-temperature string (FTS) method is utilized to identify the minimum free-energy pathways (MFEPs)[19, 20]. Specifically, we employ collective variable (CV) based string methods which utilize translationally and rotationally invariant descriptors, effectively decoupling global motions from internal conformational changes compared to Cartesian coordinates based string methods. This formulation enables the string to capture cooperative motions that are difficult to resolve in Cartesian space, such as collective solvent rearrangements, making it highly suitable for the purpose of this work.

We select physically motivated CVs to parameterize the string and elucidate distinct permeation mechanisms. As illustrated in Figure 2, these CVs capture different physical processes directly or indirectly coupled to the ion permeation process. Specifically, the string is parameterized using the following three CVs:

1. Cation Position along the Pore Axis (*d_z_*): Defined as *d_z_* = *z*_cation_ − *z*_center_, where *z*_cation_ represents the *z*-coordinate of the permeating cation and *z*_center_ denotes the geometric center of the C*_α_* atoms of MERS ETM. This variable provides a direct measurement of the progress along the permeation axis.
2. Cation–Counterion Distance (*r*_ion_): As discussed below, counterion may participate in the cation permeation process in distinct manners. We define a smooth-minimum function as 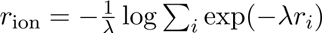, where *r_i_* is the distance between the cation and the *i*-th selected counterion, and *λ* is a smoothing parameter ensuring the variable remains continuous and differentiable. This CV monitors counterion dynamics to distinguish among three distinct transport pathways: a cation-only mechanism involving no counterion, a contact ion-pair (CIP) mechanism where a stable ion pair persists throughout transport, and a “chaperoned” ion-pair mechanism characterized by a transient ion-pair formation followed by dissociation within the pore. A similar mechanism has been discussed for multivalent-ion transfer across water-oil interface[21–23].
3. Water-Wire Connectivity (*ϕ*_ow_)[24]: This metric quantifies the structural continuity of the single-file water wire extending from the N-terminus to the C-terminus of the channel, where values approaching 1 denote an uninterrupted, full-length water wire. The formal mathematical definition of *ϕ*_ow_ is provided in the SI Appendix. This CV characterizes the global pore hydration state, which is crucial because permeating cations drag localized solvation shells into the hydrophobic pore lumen. Consequently, the instantaneous hydration state of the pore modulates the desolvation penalty, directly shaping the underlying free-energy profile.

**Figure 2:**
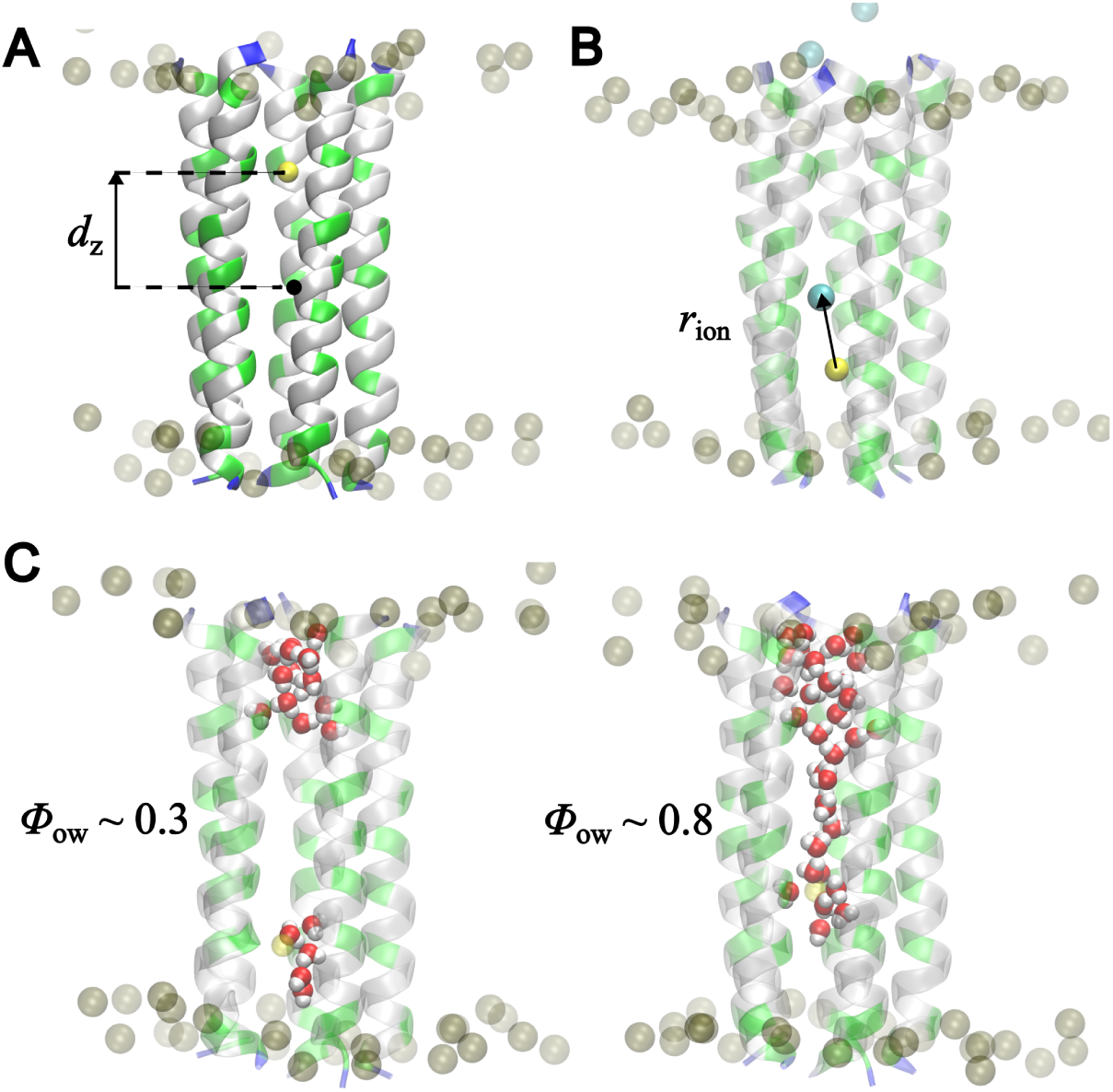
Definition of collective variables utilized in finite-temperature string simulations for the cation permeation process. (A) Position of the cation (*d_z_*) inside the MERS ETM pore relative to the pore center. (B) Distance (*r*_ion_) between the cation and the nearest counterion. (C) Water-wire connectivity (*ϕ*_ow_) from the N-terminal to the C-terminal of the MERS ETM, illustrated by two representative snapshots displaying low and high (*ϕ*_ow_) values. The permeating Na^+^ cation and its accompanying counterion Cl*^−^* (see text) are displayed as yellow and cyan spheres, respectively. The boundaries of the lipid bilayer are displayed by the phosphorus (P) atoms of the lipid headgroups, rendered as brown spheres.

Beyond path parameterization, the accuracy of string simulations depends critically on the description of underlying interactions. Ions experience electronic screening and dielectric responses from local water molecules and neighboring protein residues during permeation through the MERS ETM. Traditional nonpolarizable force fields utilize fixed point charges and fail to capture many-body polarization effects, often artificially overestimating desolvation penalties and free energy barriers[25, 26]. Incorporating a polarizable force field[27–30] is therefore potentially critical to an accurate description of the energetics of the permeation pathway. In this work, we explicitly compare the CHARMM36m[31] and the polarizable Drude[30] force fields to systematically evaluate the impact of polarization on ion permeation.

### 2.2 Free Energy Landscapes and Minimum Free Energy Pathways with the CHARMM36m Force Field

Figure 3 illustrates the MFEPs and the corresponding potential of mean force (PMF) for sodium cation (Na^+^) permeation through the MERS ETM, as resolved by FTS simulations. As described in the previous section, the pathway is parameterized via a three-dimensional CV space (*d_z_*, *r*_ion_, *ϕ*_ow_). The permeation process initiates at the reactant state (Image 0), where the Na^+^ is located above the N-terminal pore of the channel. This equilibrated initial configuration is characterized by *d_z_* ≈ −30 Å, a *ϕ*_ow_ value of approximately 0.3, and an *r*_ion_ range of 3 Å to 6 Å, corresponding to a mixture of CIP and solvent-separated ion pairs (SSIP). The transition terminates at the product state (Image 29), where the cation has migrated past the C-terminal exit. Here, *d_z_* ≈ 30 Å, *ϕ*_ow_ returns to a baseline level comparable to the initial state, and *r*_ion_ varies depending on the specific counterion participation mechanism.

**Figure 3:**
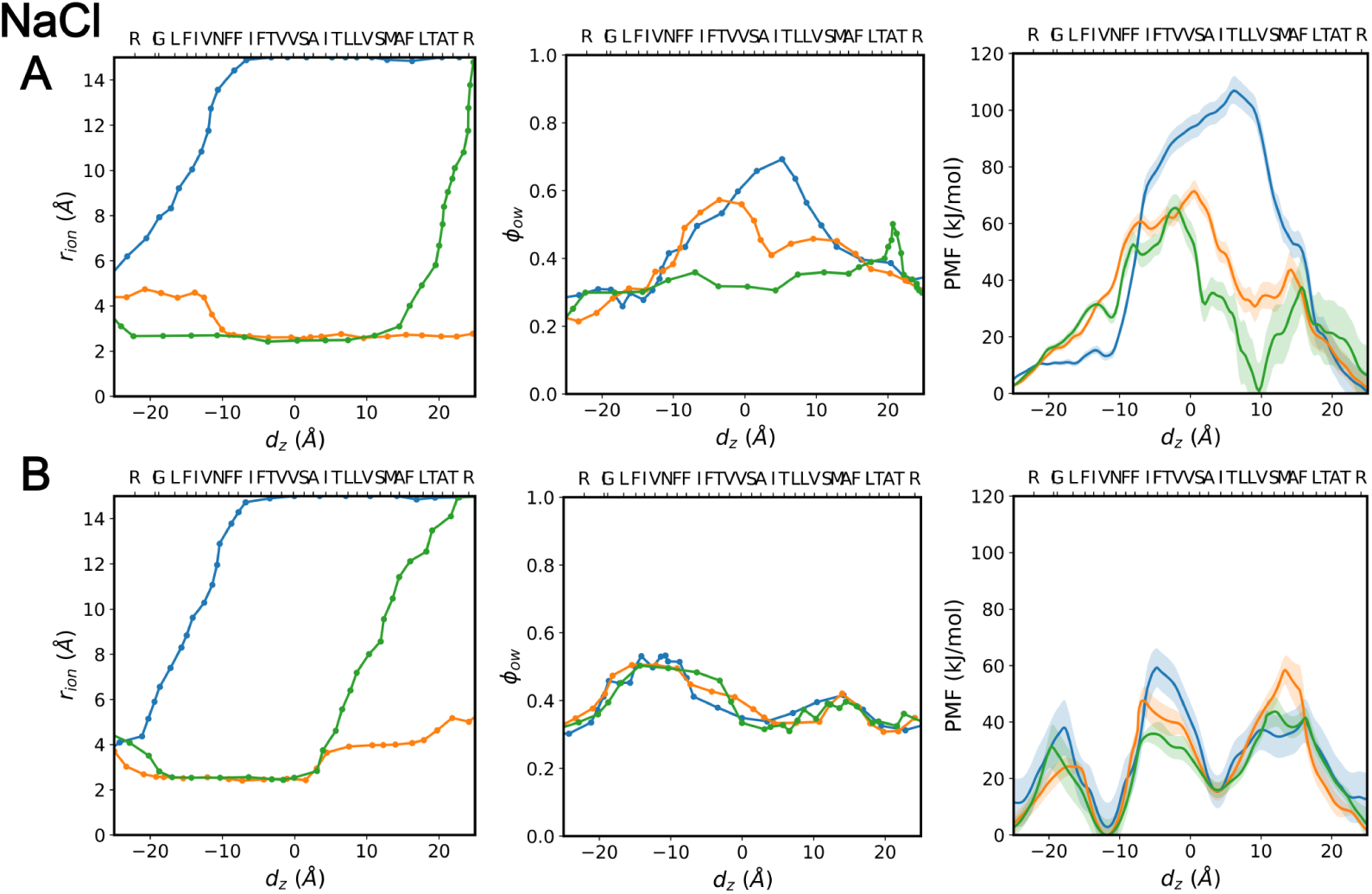
Minimum free energy pathways (MFEPs) and corresponding potentials of mean force (PMFs) for Na^+^ permeation through the MERS ETM, with Cl*^−^* as counterion. Results are shown in (A) CHARMM36m and (B) polarizable Drude force fields. Individual columns display the minimum free energy pathways projected onto the (*d_z_*, *r*_ion_) and (*d_z_*, *ϕ*_ow_) subspaces, alongside the corresponding PMFs reweighted along *d_z_*. Profiles are presented for the three distinct mechanisms: cation-only (blue), contact ion pair (orange), and chaperoned ion pair (green). MERS ETM residue locations along the channel axis are indicated by their respective C*_α_* positions.

As the system evolves along the MFEP, simulations utilizing the CHARMM36m force field (Figure 3A) reveal distinct features across the different cation permeation mechanisms. For the cation-only mechanism, *r*_ion_ rapidly increases to approximately 15 Å as the cation traverses from *d_z_* = −20 Å to −10 Å. This region corresponds to the N-terminal pore segment surrounded by residues G10 to F16, across which the PMF profile remains relatively flat. Concurrently, *ϕ*_ow_ remains constant up to *d_z_* ≈ −10 Å before steadily rising to a local maximum of *ϕ*_ow_ ≈ 0.7 at *d_z_* ≈ 4 Å (Image 19), which structurally correlates with the position of Na^+^ near residue T26. Notably, this configuration coincides with the global free-energy maximum, presenting an activation barrier of approximately 110 kJ/mol. Consequently, Image 19 (*d_z_* ≈ 4 Å, *r*_ion_ ≈ 15 Å, and *ϕ*_ow_ ≈ 0.7) represents the transition state (TS) for the cation-only permeation mechanism. Beyond this activation barrier, the system relaxes into the product basin at Image 29 to complete the transition with rather steep and monotonic decrease in the free energy.

These thermodynamic and structural profiles demonstrate that the free-energy barrier for ion permeation is strongly coupled to the pore hydration state. This coupling is evident from the flat PMF in the region of *d_z_* = −20 Å to −10 Å, where *ϕ*_ow_ maintains a baseline value of 0.3; the free energy begins to rise only when the water wire expands in concert with the advancing cation. The highly synchronous behavior between *d_z_* and *ϕ*_ow_ indicates that pore hydration and ion permeation operate via a highly cooperative, rather than a simple stepwise, mechanism.

For the ion-pair-mediated permeation mechanism, the MFEP exhibits significant differences compared to the cation-only mechanism. Upon entry of the Na^+^ into the pore, *r*_ion_ decreases from approximately 5 Å to 3 Å, indicating a transition from a solvent-separated ion pair to a contact ion pair. A significant difference is observed in the water-wire connectivity; *ϕ*_ow_ remains below 0.6 across the entire MFEP, whereas it reaches a maximum of 0.7 during cation-only permeation. This suppression indicates that ion-pair-mediated permeation requires less extensive pore hydration. The co-permeating counterion provides electrostatic screening for the cation, thereby mitigating the desolvation penalty without inducing the thermodynamically unfavorable insertion of additional water molecules into the highly hydrophobic pore. This mechanism is energetically supported by the computed potential of mean force, which reveals a global activation barrier of approximately 70 kJ/mol, roughly 40 kJ/mol lower than that calculated for the cation-only pathway.

Furthermore, the free-energy landscape for the ion-pair-mediated mechanism is more rugged, featuring two distinct activation barriers located at *d_z_* ≈ 0 Å (near residue A21) and *d_z_* ≈ 15 Å (near residue F33). A metastable intermediate basin is resolved at *d_z_* ≈ 10 Å, which corresponds to the region bounded by the polar residues T26 and S30 and suggests a transient cation-binding site within the pore.

A contact ion-pair mechanism where the pair of ions remain associated throughout the permeation process is incompatible with electrophysiological data. The translocation of a net-neutral, dipolar, species would yield zero electrical conductance, directly contradicting the experimentally measured wild-type MERS E protein channel conductance of 113.5 ± 12.5 pS[14]. By contrast, the “chaperoned” ion-pair mechanism features a stable ion pair as the cation enters and traverses the majority of the pore, followed by dissociation at *d_z_* ≈ 12 Å near the F33 residue. Here, the Na^+^ passes through the F33 gate while the counterion Cl*^−^* exits the channel in the opposite direction to return to the original side of the membrane. This dissociation and exit event demands extra channel hydration, as evidenced by the late peak in *ϕ*_ow_ at *d_z_* ≈ 20 Å. Because it facilitates the translocation of a net charge, the chaperoned ion-pair mechanism qualitatively aligns with experimental conductance observations. However, the calculated free-energy barrier (∼60 kJ/mol) remains significantly higher than the first-order, one-dimensional electrodiffusion estimate of a few *k_B_T* inferred from macroscopic conductance (see SI Appendix). The quantitative discrepancy, coupled with the tight correlation between *ϕ*_ow_ and the PMF profile, strongly suggests that the non-polarizable CHARMM36m force field fails to accurately capture the pore hydration state and ion stabilization during ion permeation, resulting in an significantly elevated free-energy barrier.

### 2.3 Free Energy Landscapes and Minimum Free Energy Pathways with the Drude Force Field

The Drude polarizable force field reveals substantially different molecular features for all three permeation mechanisms compared to the CHARMM36m model. While the evolution of *r*_ion_ follows a similar trajectory for the cation-only and CIP mechanisms, a notable divergence occurs in the chaperoned ion-pair mechanism: dissociation of the ion pair occurs earlier at *d_z_* ≈ 4 Å near residue T26 rather than near residue F33 (see additional discussions below). The most significant difference between the two force fields lies in the profile of *ϕ*_ow_. In contrast to the pathway-dependent *ϕ*_ow_ behaviors observed in CHARMM36m, the Drude model yields remarkably uniform *ϕ*_ow_ trends across all three permeation pathways. Specifically, *ϕ*_ow_ begins to rise at *d_z_* ≈ −20 Å, reaching an initial local maximum at *d_z_* ≈ −10 Å near residue N15. Subsequently, *ϕ*_ow_ decreases to a local minimum at *d_z_* ≈ 4 Å near residue T26, before rising to a second local maximum at *d_z_* ≈ 12 Å in the vicinity of residue F33. Finally, the system relaxes into the product basin at the terminal image, with *ϕ*_ow_ returning to its baseline value of approximately 0.3, completing the transition.

Similar to the CHARMM36m results, the PMF profiles generated by the Drude simulations remain strongly coupled to the evolution of *ϕ*_ow_. This persistent correlation underscores that the pore hydration state remains the decisive coordinate that governs the permeation process. Consequently, because the underlying hydration profiles are uniform across the three mechanisms, the PMF profiles also converge onto a nearly identical shape. Crucially, the global free-energy maximum decreases dramatically to approximately 60 kJ/mol for the cation-only pathway and 40 kJ/mol for the chaperoned ion-pair pathway. Compared to the corresponding CHARMM36m values of 110 kJ/mol and 70 kJ/mol, respectively, this ∼ 50% reduction in the activation energy represents a significant improvement, bringing the calculated thermodynamic barriers into a realistic range.

A simple physical explanation for this attenuated free-energy barrier can be developed by considering electronic polarization and its influence on the local dielectric environment of the permeating ion(pair). According to the Born model of ion solvation[32], the desolvation penalty of a spherical ion is directly proportional to the dielectric difference term Δ(1*/ɛ_r_*) = (1*/ɛ*_pore_ − 1*/ɛ*_bulk_). In the CHARMM36m model, the fixed point charges limit the empty pore to an unpolarized vacuum state where *ɛ*_pore_ ≈ 1, leading to an abrupt change in dielectric environment as the ion transits from bulk water (*ɛ*_bulk_ ≈ 80) to the pore, leading to a significant desolvation penalty. By contrast, the Drude model captures the electronic response of the hydrophobic channel by oscillating Drude particles, correctly accommodating a higher dielectric constant of *ɛ*_pore_ ≈ 2. Evaluating the Born expression under these conditions reveals that the desolvation penalty is scaled by a factor of ∼ 0.5 when explicit polarization are included. This extra dielectric screening partially accounts for the approximately twofold reduction in the computed free-energy barriers with the Drude model.

We also analyzed the gating effects of the N-terminal (F12, F16, F17, F19) and C-terminal (F33) phenylalanine residues by calculating the local resistivity profiles (see SI, Fig. S7). The profiles reveal that while both regions show significant gating control, residue F33 exhibits a more pronounced effect, with a local resistivity an order of magnitude higher than that of the N-terminal cluster. Additional analysis across the string images with the permeating cation near residue F33 demonstrates that the polarizable Drude force field captures the cation–*π* interaction with significantly higher accuracy than CHARMM36m model (see SI Appendix, Fig. S1). Further evaluations on idealized model systems reveal that this trend remains robust across broad angular orientations as the Na^+^ approaches the phenylalanine (Phe) side chain from both vertical and lateral directions (Fig. S3). Specifically, when Na^+^ approaches the Phe side chain vertically along the ring normal, the Drude force field successfully reproduces the reliable DFT equilibrium distance within ∼ 0.2 Å and the target interaction energy within an error of ∼ 10 kJ/mol; the error in CHARMM36m is, by comparison, ∼ 45 kJ/mol averaged over the snapshots and can be larger than 120 kJ/mol for some configurations. As the approach direction changes from vertical to horizontal, the Drude model systematically overestimates the cation–*π* interaction energy, while CHARMM36 predicts the interaction to be purely repulsive, in contrast to DFT (see additional discussions in the SI) and higher-level quantum calculations[33, 34]. However, the overpolarized configurations are rarely sampled in the FTS simulations (Fig. S2B) and therefore do not impact the accuracy of the Drude simulations for ion permeation.

A detailed inspection of the Drude PMF features reveals several distinct transition states and metastable intermediates along the MFEP. Here, we analyze the trajectory corresponding to the chaperoned ion-pair mechanism, as it represents the most energetically favorable pathway and explicitly captures the coupling among all primary coordinates. Along this profile, we identify seven key configurations of interest whose structural snapshots are displayed in Figure 4, labeled panels (A) through (G).:

A. The approach of the cation toward the N-terminal pore (*d_z_* ≈ −25 Å);
B. The first transition state (TS_1_ at *d_z_* ≈ −17 Å), localized near residue F12;
C. The first intermediate (Int_1_ at *d_z_* ≈ −11 Å), localized near residue N15;
D. The second transition state (TS_2_ at *d_z_* ≈ −5 Å), localized near residue F19;
E. The second intermediate (Int_2_ at *d_z_* ≈ 4 Å), localized near residue T26;
F. The third transition state (TS_3_ at *d_z_* ≈ 12 Å), localized near residue F33;
G. The exit of the cation past the C-terminal gate (*d_z_* ≈ 25 Å);

**Figure 4:**
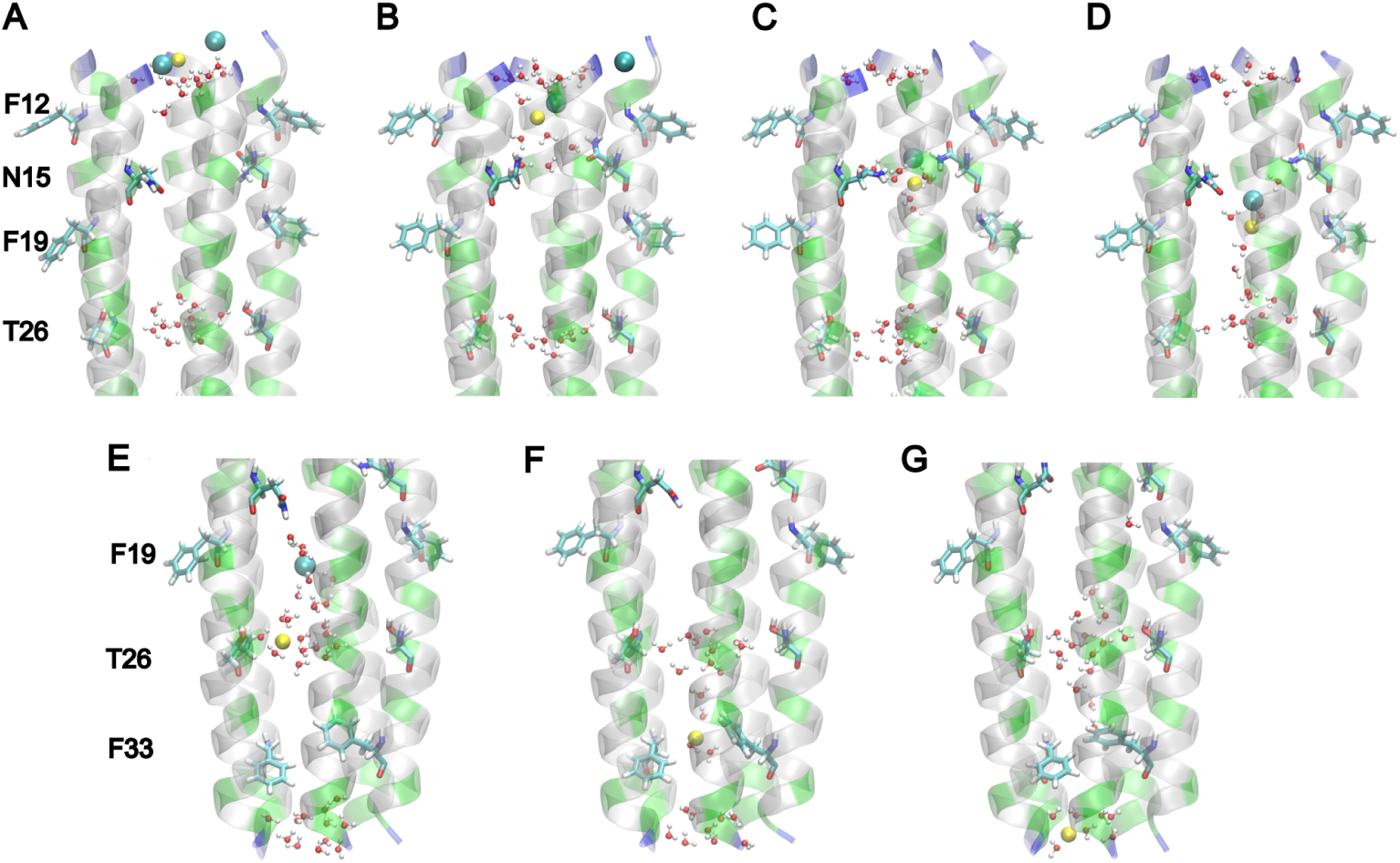
Representative configurations of Na^+^ permeation through the MERS ETM pore via the chaperoned ion-pair mechanism. Snapshots were obtained using the polarizable Drude force field. The permeating Na^+^ cation and its accompanying counterion Cl*^−^* are displayed as yellow and cyan spheres, respectively. Sequential panels depict key states along the pathway: (A) reactant state, (B) transition state 1 (TS_1_), (C) intermediate 1 (Int_1_), (D) transition state 2 (TS_2_), (E) intermediate 2 (Int_2_), (F) transition state 3 (TS_3_), and (G) product state. Critical channel residues, including aromatic residues (F12, F19, F33) and polar residues (N15, T26), are highlighted using the same color scheme as in Figure 1 (polar in green, hydrophobic in gray). An expanded view emphasizing the local interactions of the permeating ion with neighboring water molecules and protein residues is provided in SI Appendix (Fig. S6).

Mechanistically, the three activation barriers are tightly coupled to the localized water wires across hydrophobic pore segments that lack baseline hydration in the resting state. At TS_1_, a localized water wire bridges the N-terminal entrance to residue N15. At TS_2_, a continuous water wire spans the region between residues N15 and T26. Finally, TS_3_ involves the extension of the water wire from T26 through the hydrophobic F33 constriction to the C-terminal exit. Notably, the structural solvation shell around residue T26 must be partially disrupted to accommodate the long-range reorganization required for water-wire development during both the TS_2_ and TS_3_ transitions.

The two intermediate basins coincide with polar residues that exhibit either structural or state-dependent transient solvation. During the occupancy of Int_1_, the side chain of N15 which remains dry in the equilibrium resting state undergoes transient wetting as the ion-pair traverses the region; this is in agreement with recent ssNMR study that observed locally enhanced solvation near N15/F17 upon K^+^ binding[18]. At Int_2_, the side chain of T26 possesses a structurally stable solvation shell that accommodates the permeating cation, which has fully dissociated from its counterion by this stage.

### 2.4 Changes in Local Hydration States During Ion Permeation with Polarizable Drude Models

By definition, the *ϕ*_ow_ quantifies the continuous structural connectivity of the water wire from the N-terminus to the C-terminus. However, as demonstrated in the preceding section, the formation of water wires often involves the structural reorganization of localized solvation shells, to which a global CV like *ϕ*_ow_ is inherently less sensitive. To monitor these localized hydration dynamics with higher spatial resolution, we evaluate the local water molecule number density profiles around individual residues as a function of the reaction progress, parameterized by the string simulation image index (*s*) ranging from 0 to 30, as illustrated in Figure 5. Integrating the thermodynamic profiles from the MFEP and PMF with these localized hydration landscapes allows for a rigorous evaluation of pore hydration states and the identification of intermediate basins acting as functional cation-binding sites.

**Figure 5:**
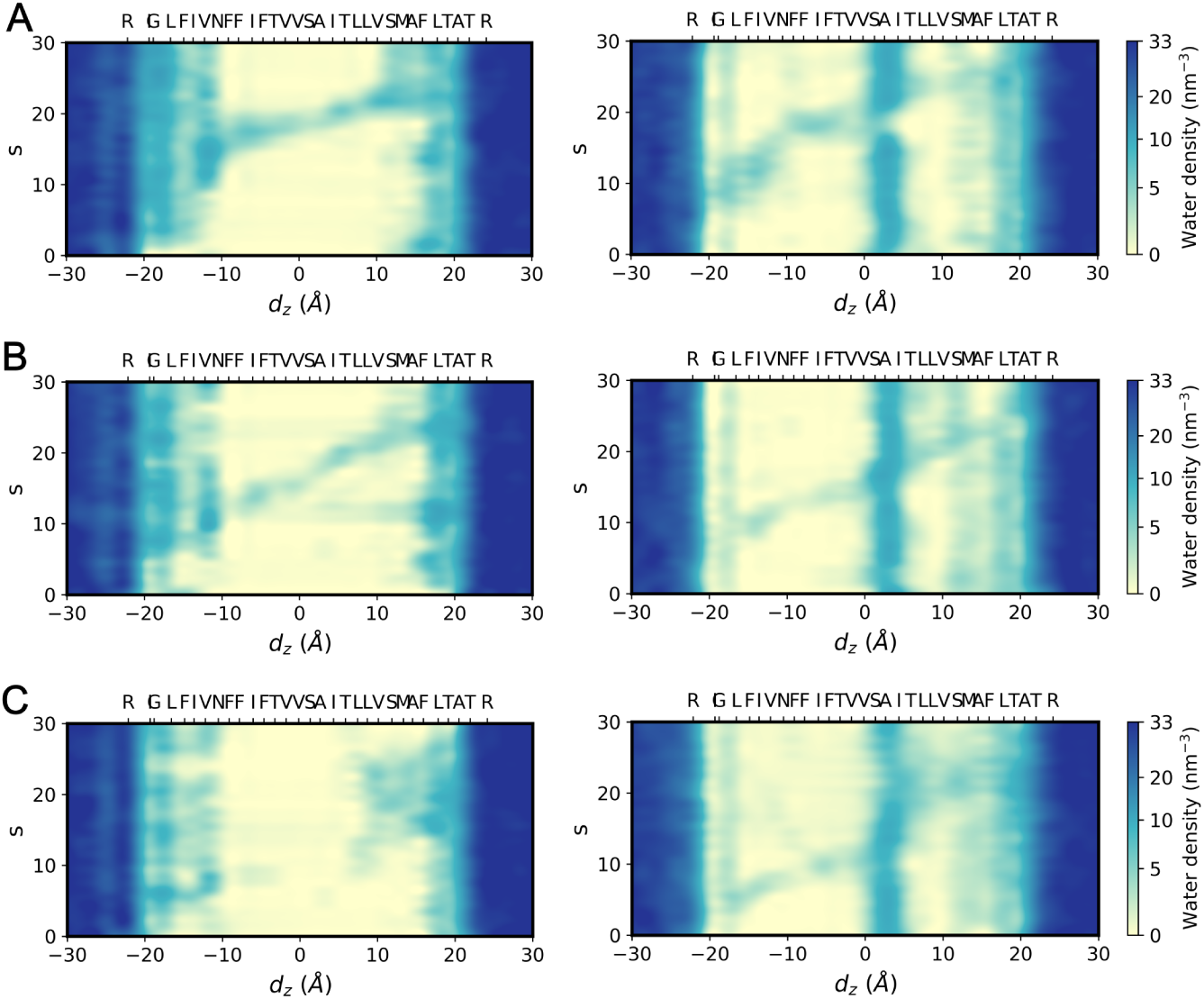
Water number density (in nm*^−^*^3^) profiles along the pore axis during the Na^+^ permeation process. Profiles are plotted as a function of (*d_z_*) and reaction coordinate (*s*) defined from the images used in finitetemperature string simulations for the three mechanisms: (A) cation-only, (B) contact ion-pair, and (C) chaperoned ion-pair. Results obtained using the additive CHARMM36m and polarizable Drude force fields are displayed in the left and right columns, respectively. MERS ETM residue locations along the channel coordinate are indicated by their respective C*_α_* carbon positions.

Across the localized hydration profiles of all three ion permeation pathways, the equilibrium baseline hydration of the pore exhibits significant differences between CHARMM36m force field and the polarizable Drude model. In CHARMM36m, baseline hydration is primarily distributed near the N-terminal region above residue N15 and the C-terminal region below residue F33; the central pore lumen remains completely dry, devoid of persistent water molecules. By contrast, the Drude model yields a remarkably different distribution: the N-terminal region between F12 and N15 remains mostly dewetted in the equilibrium resting state while a stable, structural solvation shell centers around the polar residues S23 and T26 within the mid-pore lumen, as shown in Figure 5. Furthermore, the Drude PMF (Figure 3) and the intermediate configuration Int_1_ (Figure 4C)demonstrate that N15 undergoes induced, transient solvation in the presence of proximal ions. Given that the relative free energy of this intermediate state is approximately the same as the bulk solution basin, residue N15 can functionally serve as a localized cation-binding site near the N-terminal pore. The difference in pore hydration state between CHARMM36m and Drude can be explained by the introduction of explicit polarization. Hydrogen bonding between C=O and N-H groups within the *α*-helix induces a local dipole enhancement that is captured by the polarizable Drude force field, but absent in the fixed-charge CHARMM36m model. The resulting localized polarization makes the inner pore wall and specific polar residues more hydrophilic, thereby stabilizing the local confined water networks. Polarization of nonpolar residues also contributes to the effective pore dielectric, although its role in stabilizing the local hydration is minor, as theses interactions are primarily due to transient induced dipoles rather than the stronger, permanent dipoles from polar residues.

Consequently, the stably solvated region surrounding the polar residues S23/T26 acts as a thermodynamic relay location for ion permeation. Within the framework of the CHARMM36m model, the absence of this intrinsic mid-pore hydration forces the permeating ion to extensively alter the pore environment by dragging a massive number of water into an otherwise hydrophobic lumen. This high desolvation penalty applies to both the cation-only and the ion-pair-mediated mechanisms; as shown in the left panels of Figure 5, ion permeation requires hydration of either the entire channel or the substantial segment spanning from T26 to the C-terminus, both of which are thermodynamically prohibitive. By contrast, the Drude model leverages the intrinsic baseline hydration at S23/T26 and the transiently induced hydration at N15 to establish a series of spatial relays, as shown in the right panels of Figure 5. Rather than inducing a large-scale, unfavorable flooding of the channel, ion permeation is achieved via the sequential formation of short, localized water wires between these relay locations. This mechanism minimizes disruption to the global channel hydration state, yielding a highly favorable free-energy pathway.

Under this polarizable framework, the co-permeating counterion provides essential electrostatic screening, allowing the cation to more easily traverse the initial dewetted segment from the N-terminus to N15, and subsequently to T26, thereby lowering the initial activation barriers. Upon reaching the stable hydration pocket at residue T26, the ion pair undergoes dissociation, diverging from the late-stage dissociation near F33 observed in CHARMM36m simulations. This early separation allows the counterion to exit the pore along a significantly shortened path, minimizing further disruption to the pore hydration state. Collectively, these cooperative electrostatic and hydration phenomena explain why the global free-energy barrier for ion permeation is dramatically attenuated in the polarizable Drude model compared to CHARMM36m, especially in the chaperoned ion pair mechanism, bringing the computational findings into closer alignment with experimental electrophysiological data.

### 2.5 Impact of Counterion Substitution as a Mechanistic Probe

With the counterion mediated mechanisms, the identity of the counterion is expected to have an impact on the permeation process, providing a potential experimental probe of these pathways. Accordingly, we performed string simulations substituting Cl*^−^* with Br*^−^*, while maintaining other parameters identical across both the CHARMM36m and Drude force fields. As illustrated in Figure 6, the resulting MFEPs and corresponding PMF profiles display overall similar features compared with those obtained from the Cl*^−^* simulations across all three permeation pathways. In the non-polarizable CHARMM36m simulations, the cation-only pathway exhibits a transition state localized at *d_z_* ≈ 4 Å (near residue T26) with an activation barrier of approximately 100 kJ/mol, while both CIP and chaperoned ion-pair mechanisms yield free-energy barriers spanning 70 to 80 kJ/mol. In the Drude polarizable simulations, the rugged feature of the free-energy landscape defined by three distinct transition states and two metastable intermediates (localized at residues N15 and T26) is preserved. These observations demonstrate that the underlying molecular features of the three permeation mechanisms remain robust against counterion substitution.

**Figure 6:**
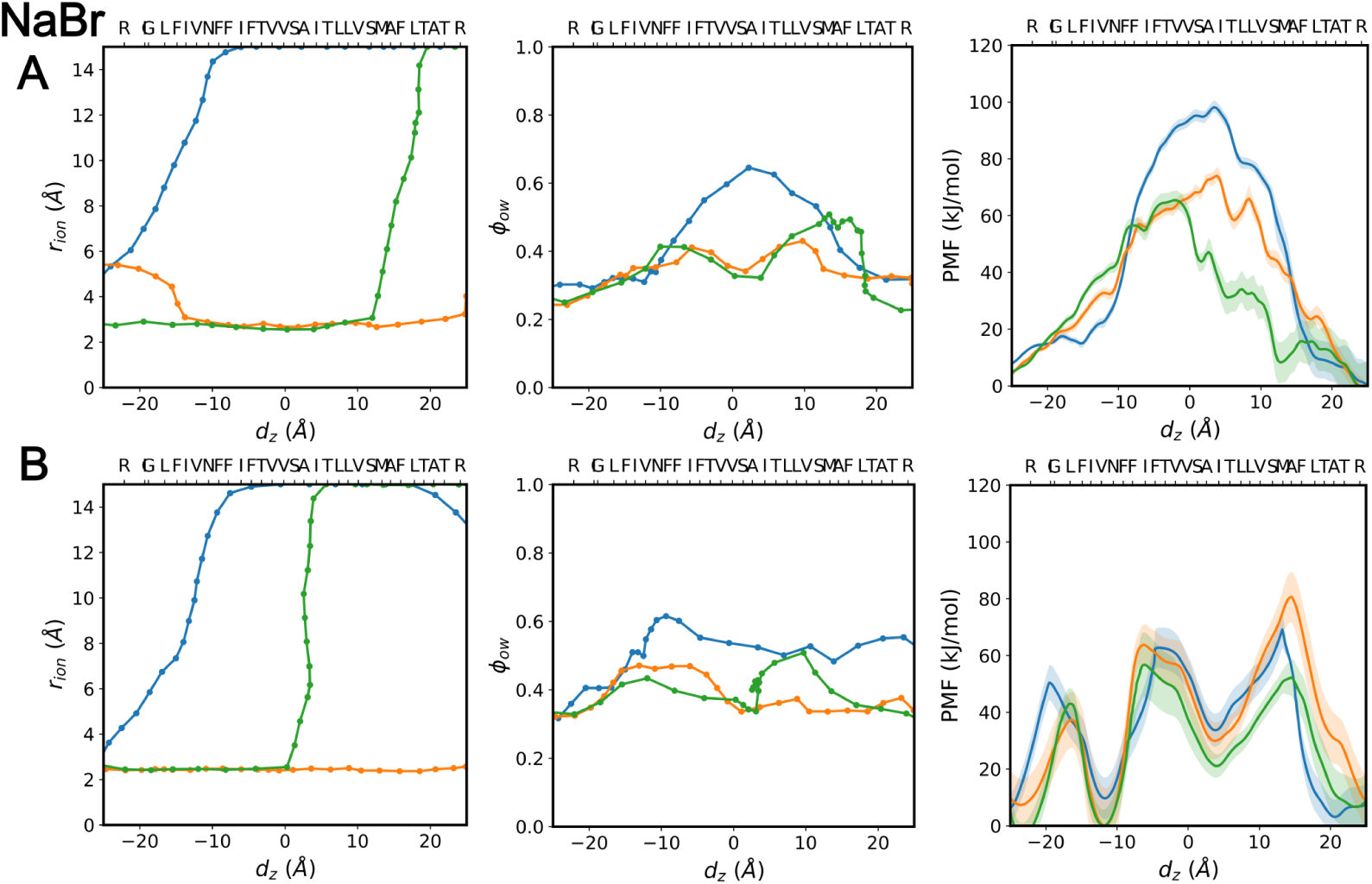
Minimum free energy pathways (MFEPs) and corresponding potentials of mean force (PMFs) for Na^+^ permeation through the MERS ETM, with Br*^−^* as counterion. Results are shown in (A) CHARMM36m and (B) polarizable Drude force fields. Individual columns display the minimum free energy pathways projected onto the (*d_z_*, *r*_ion_) and (*d_z_*, *ϕ*_ow_) subspaces, alongside the corresponding PMFs reweighted along *d_z_*. Profiles are presented for the three distinct mechanisms: cation-only (blue), contact ion pair (orange), and chaperoned ion pair (green). MERS ETM residue locations along the channel axis are indicated by their respective C*_α_* positions.

Importantly, a quantitative difference emerges within the polarizable framework: for the ion-pair-mediated mechanisms, the Drude simulations reveal an elevated free-energy barrier for the Br*^−^* counterion system compared to its Cl*^−^* counterpart (60 kJ/mol versus 40 kJ/mol). If the physiological transport process is predominantly driven by an ion-pair-mediated mechanism, this calculated thermodynamic penalty suggests that substituting chloride for bromide in the background electrolyte would increase the global activation barrier, thereby systematically depressing single-channel conductance. This computationally derived trend serves as a testable hypothesis for future electrophysiological studies, offering a clear strategy to experimentally probe the existence of the chaperoned ion-pair mechanism and to further elucidate the conduction mechanism of the MERS E and related proteins.

## 3 Concluding Remarks

The molecular mechanism by which ions traverse predominantly hydrophobic pores has remained an important open question because the energetic penalty associated with ion desolvation appears fundamentally incompatible with efficient conduction. Indeed, dewetting transition associated with hydrophobic cavities has been discussed as a gating mechanism that blocks ion transport in more canonical ion channels[6–10]. Using the MERS coronavirus E protein as a model system, our simulations provide a plausible resolution of this apparent paradox. Rather than crossing the pore as an isolated hydrated ion, the permeating cation is assisted by a cooperative process involving transient counterion association, localized hydration relays, and electronic polarization. Together, these effects substantially reduce the desolvation penalty while preserving efficient charge transport through an otherwise unfavorable hydrophobic environment.

Among these factors, the active participation of counterions represents the most conceptually unexpected finding. Although transient ion pairing has previously been discussed for transport across liquid–liquid interfaces [21, 22], counterions have generally been regarded as passive components of the electrolyte background in biological ion transport. Our analyses instead suggest that counterions transiently become part of the permeation process, screening electrostatic interactions at the most unfavorable stages of transport before dissociating to preserve net ionic conduction. This mechanism explains why ion permeation can occur without extensive wetting of the hydrophobic pore.

Equally importantly, our results demonstrate that pore hydration does not merely transition between dry and wet states as previously discussed in channels[6, 7, 35] and transporters/pumps[24, 36–40]. Instead, efficient conduction emerges through the sequential formation of localized “hydration relays” that transiently connect pre-existing hydration sites as the ion progresses through the channel. This dynamic coupling between ion position, local hydration, and electrostatic screening defines the underlying free-energy landscape and provides a more nuanced picture of hydration-assisted transport than a simple wetting transition.

Our study also highlights the critical importance of explicitly accounting for electronic polarization when modeling ion transport through heterogeneous environments [10, 41–44]. Beyond quantitatively lowering the free-energy barrier, polarization qualitatively alters the hydration landscape, stabilizes cation–*π* interactions, and changes the preferred permeation mechanism. These observations suggest that additive force fields may systematically overestimate desolvation barriers in hydrophobic pores and underscore the importance of polarizable models for quantitative studies of viroporins and related membrane channels.

The calculated free-energy barriers remain several kcal/mol higher than estimates derived from experimental conductance. Consequently, applying an inhomogeneous diffusion model to the simulated PMF and position-dependent diffusion coefficients yields a conductance several orders of magnitude lower than experimental measurements (see SI). As discussed in recent ion channel literature[44–46], predicting absolute conductance remains challenging due to uncertainty in the available structural models and biomolecular force fields[45, 46], thus the discrepancy most likely reflects these technical limitations rather than the mechanistic picture emerged from our analysis. The pore radius of the structural model used here is rather narrow (Figure 1B), and recent ssNMR study[18] showed that K^+^ binding increases the structural homogeneity of the MERS ETM, although without major structural transitions. Moreover, we choose Na^+^ to establish a baseline of electronic polarization during cation permeation through a hydrophobic pore. Due to its high charge density, Na^+^ enforces a tightly ordered first solvation shell that imposes a large penalty on desolvation and water-wire reorganization. Substituting Na^+^ with K^+^, which possesses a lower charge density and a more flexible and loosely bound solvation shell, should further reduce the desolvation and water-wire disruption penalties. Collectively, these effects are expected to shift the free-energy landscape toward a lower-barrier permeation profile. Future work integrating improved structural ensembles, focusing specifically on the conformations of the cation-binding site near residue N15 and the gating phenylalanine residues (particularly F33), with nonequilibrium simulations under applied membrane potentials should enable increasingly quantitative descriptions of viral ion-channel conductance.

More broadly, the physical principles identified here are unlikely to be unique to the MERS E protein. Cooperative electrostatic screening, localized hydration relays, and electronic polarization may represent a general strategy by which biological and synthetic hydrophobic nanopores overcome the fundamental thermodynamic penalty associated with ion desolvation. We anticipate that these concepts will aid both the mechanistic interpretation of ion transport in membrane proteins and the rational design of artificial nanopores with tailored transport properties.

## 4 Materials and Methods

### 4.1 Preparation Protocols for CHARMM36m and Drude Simulation Systems

The initial coordinates of the pentameric channel were derived from the solid-state NMR structure of the MERS ETM (PDB ID: 9DCJ[14]), retaining residues 8–38 for the simulation models. All simulation systems were constructed using the CHARMM-GUI interface[47]. To closely mimic the lipid environment of the ERGIC, a symmetric membrane bilayer was generated; each leaflet was composed of 39 1-palmitoyl-2-oleoylsn-glycero-3-phosphocholine (POPC), 12 1-palmitoyl-2-oleoyl-sn-glycero-3-phosphoglycerol (POPG), and 9 cholesterol (CHOL) molecules, yielding a final molar ratio of 65:20:15 (POPC:POPG:CHOL). The bilayer was solvated with a 4-nm-thick water layer on each side, producing a final simulation box with approximate dimensions of 6 × 6 × 13 nm^3^. The aqueous phase was neutralized and maintained at a physiological concentration of 150 mM NaCl by adding 48 Na^+^ and 34 Cl*^−^* ions. The protein and lipids were parameterized using the CHARMM36m[31] force field, water molecules were modeled via the TIP3P[48] formulation, and ions were treated using the Joung-Cheatham 12-4 parameters[49].

Energy minimization and subsequent equilibration workflows were executed using the GROMACS 2021[50, 51] software package. The system initially underwent a 100-ns stepwise equilibration protocol within the canonical (NVT) ensemble, partitioned into eight consecutive 10-ns stages followed by a final 20-ns equilibration phase. Throughout the first eight stages, harmonic position restraints were applied to the protein backbone atoms and lipid heavy atoms, with the corresponding force constants systematically scaled down from 400 to 200, 50, 20, 10, 5, 2, and 1 kJ*/*(mol · Å^2^). During the final 20-ns equilibration stage, all harmonic position restraints were released to allow unconstrained relaxation of the ETM. The system temperature was regulated at 298 K using a stochastic velocity-rescaling thermostat[52]. Short-range, real-space electrostatic and van der Waals interactions were truncated at a cutoff distance of 1.2 nm, and long-range electrostatic interactions were evaluated using the Particle Mesh Ewald (PME) method[53]. All covalent bonds involving hydrogen atoms were constrained via the LINCS algorithm [54], enabling the utilization of a standard 2-fs integration time step across all production trajectories.

To investigate the impact of explicit electronic polarization, the final configuration of the MERS ETM obtained from the 100-ns equilibration phase served as the starting configuration for subsequent polarizable production runs. These simulations were parameterized using the polarizable Drude 2023 force field [30]. Due to the absence of published parameterization for POPG and cholesterol within the Drude framework, a pure POPC lipid bilayer of comparable dimensions was substituted. To prevent non-physical structural drift of the pentameric bundle in this simplified membrane environment, a weak harmonic position restraint of 10 kJ*/*(mol · Å^2^) was applied to the protein backbone heavy atoms throughout the production phases. The system temperature was regulated at 298K for real atoms and 1K for drude particles using an extended-Langevin thermostat[55, 56]. All Drude polarizable trajectories were integrated with a time step of 1 fs using the OpenMM software package[57, 58].

### 4.2 Finite-Temperature String Simulation Protocols

To generate the initial guess for the finite temperature string calculations, constant-velocity steered molecular dynamics (SMD) simulations were executed across both the CHARMM36m and Drude configurations utilizing identical biasing parameters. Using the equilibrated 100-ns endpoints as starting structures, a 1.2-ns SMD trajectory was generated for each system. A Na^+^ located beneath the N-terminal pore was designated as the steered particle. The steering vector was aligned along the positive z-axis of the simulation cell, reflecting the physiological transport axis from the N-terminus to the C-terminus of the channel. The constant pull velocity was set to 0.005 nm/ps, mediated by a spring force constant of 10 kJ*/*(mol · Å^2^). To effectively sample the distinct ion-pair-mediated transport mechanisms, a secondary set of harmonic boundary walls was enforced on *r*_ion_ using the PLUMED plugin [59] during the SMD pulling phases. For the strict CIP pathway, a continuous upper boundary condition (UPPER_WALL) was implemented at *r*_ion_ = 7.0 Å with a force constant of 10 kJ*/*(mol · Å^2^). For the chaperoned ion-pair mechanism, the UPPER_WALL restraint at *r*_ion_ = 7.0 Å was maintained solely during the initial 0.5 ns of the trajectory. Over the subsequent 0.2 ns, this upper boundary was smoothly deactivated while a corresponding lower boundary condition (LOWER_WALL), applying identical force parameters, was simultaneously activated.

MFEPs were optimized using FTS with the swarms-of-trajectories approach[60, 61]. Thirty initial configurations were extracted at roughly equidistant intervals along the *d_z_* collective variable from the preceding SMD trajectories to define the starting discretized images of the path. The string was parameterized in a three-dimensional CV space (*d_z_*, *r*_ion_, *ϕ*_ow_). During each iteration of the string algorithm, every image was first harmonically restrained for 100 ps to its respective target position within the CV space to ensure local relaxation. The PLUMED restraint force constants were specified as 10 kJ*/*(mol · Å^2^) for *d_z_*, 10 kJ*/*(mol · Å^2^) for *r*_ion_, and 5000 kJ/mol for *ϕ*_ow_. Following this localized equilibration phase, an ensemble of 20 independent “swarm” replicas was spawned from each image configuration by randomly re-initializing atomic velocities from the Boltzmann distribution. These replicas were subsequently integrated short-term without active CV restraints for 10 fs using a refined integration time step of 0.5 fs. The instantaneous collective variable coordinates for each replica were monitored via PLUMED, and the computed ensemble average of these displacements was utilized to propagate the string path toward the MFEP in the subsequent iteration (see SI Appendix). Path propagation and reparameterization routines were managed via in-house scripting.

### 4.3 Potential of Mean Force Calculations Along the Minimum Free Energy Pathways

Well-tempered multiple-walker metadynamics simulations[62, 63] were performed using the PLUMED plugin to construct the PMF profiles along the MFEPs. Biasing potentials were applied along the path collective variables (PathCVs) as implemented via the PATH component in PLUMED[64, 65]; a detailed mathematical definition of these collective variables is provided in the Supporting Information. Additionally, the three CVs utilized during the FTS parameterization were tracked throughout the simulations.

To maintain the sampling window relevant to the permeation process, UPPER_WALL and LOWER_WALL were enforced at *d_z_* = ±25 Å with a force constant of 10 kJ*/*(mol · Å^2^). For the production well-tempered metadynamics runs, Gaussian hills with an initial height of 1.0 kJ/mol and a width of 0.1 were deposited every 1 ps using a bias factor of 35. Enhanced sampling was achieved using ten parallel walkers initialized from distinct configurations (differing in both coordinates and velocities) and distributed with approximately equal spacing along the *d_z_* coordinate from the prior FTS trajectories. The walkers shared their accumulated bias potentials at a frequency of every 5 ps. Each walker was propagated for 300 ns, yielding a cumulative sampling time of 3 *µ*s for PMF construction. Finally, the multi-dimensional free energy surfaces were reweighted against the *d_z_* reaction coordinate following the method described in Ref.66.

## Supporting Information Appendix (SI)

Supporting Information available for additional details for the FTS simulations, estimate for the activation free energy barrier based on experimental conductance data, data for the benchmark calculations of cation-*π* interactions, data for diffusion coefficient along pore axis, and data for magnified figure on intermediate and transition states from chaperoned ion mechanism.

## Data Availability

Data supporting this study, including all FTS and WT-METAD simulation setups and CHARMM36m/Drude configurations, cation-*π* benchmarking files, and water number density and diffusion coefficient profiles, have been deposited in the Zenodo repository at https://doi.org/10.5281/zenodo.21285033.

## Acknowledgments

This work was supported by NIH grants 1R21AI194473-01A1 to Q.C. and GM159321 to M.H. The initial exploration of the Drude polarization force field simulations was supported in part by NIH grant R35GM141930 to Q.C. The computational studies were supported in part by Computational resources provided by the Boston University Shared Computing Cluster, which is administered by Boston University Research Computing Services (www.bu.edu/tech/support/research/), are greatly appreciated. Part of the computation also used the ACES GPU cluster at Texas A & M University through allocation BIO260037 from the Advanced Cyberinfrastructure Coordination Ecosystem: Services & Support (ACCESS) program, which is supported by U.S. National Science Foundation grants #2138259, #2138286, #2138307, #2137603, and #2138296.

## Supporting Information Text

### Finite-temperature String Simulation Setup

#### Cation-Counterion Distances

The smoothing factor employed in the definition of *r*_ion_ is set to be *λ* = 500. For optimal graphical visualization, the raw *r*_ion_ value was smoothly truncated using a Heaviside-conditioned switching function to yield the displayed metrics: *r*_ion, display_ = *r*_ion, raw_ · *H*(*r*_cap_ −*r*_ion, raw_)+(*r*_cap_ +tanh (*r*_ion, raw_ − *r*_cap_))· *H*(*r*_ion, raw_ −*r*_cap_), where *r*_cap_ = 14.0 Å and *H*(*x*) is the Heaviside step function. This smoothing transformation prevents visualization artifacts that arise when the cation resides deeply within the pore lumen where the raw *r*_ion_ values diverge asymptotically without corresponding to any physically meaningful ion-pair transitions.

#### Water-wire Connectivity

The water-wire connectivity parameter, *φ*_ow_, was evaluated according to equations [1]–[3]. First, a series of *N* = 28 virtual sites was defined to map the channel pore, with each site located at the geometric center of five C*_α_* atoms of the same residue index (residues Gly10 to Thr37) from each *α*-helix. The local water coordination number, *s_i_*, around the *i*-th virtual site was calculated via equation [1] using a cubic switching function:

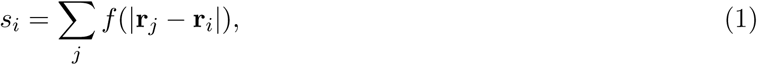

which sums over the distance between *i*-th virtual site **r***_i_* and oxygen atom from *j*-th water molecule **r***_j_*. The switching function is defined as *f* (*r*) = (*y* − 1)^2^(1 + 2*y*) for *y* ∈ (0, 1), where the normalized distance parameter *y* is defined as *y* = (*r* − *r*_1_)*/*(*r*_0_ − *r*_1_) with cutoff thresholds set to *r*_1_ = 3.0 Å and *r*_0_ = 4.5 Å. The switching function *f* (*r*) satisfies the boundary conditions *f* (*r*) = 1 for *r < r*_1_ and *f* (*r*) = 0 for *r > r*_0_, providing a smooth interpolation from 1 to 0 over the interval *r* ∈ [*r*_1_*, r*_0_].

To map these values onto a continuous probability scale, the local coordination number *s_i_*was transformed into a normalized occupancy *I_i_* using equation [2],

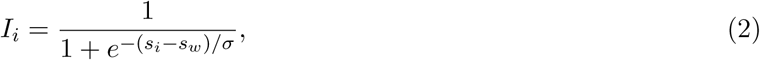

which is parameterized with a reference coordination number threshold *s_w_*= 1.25 and a smoothing factor 1*/σ* = 1.5.

Finally, the comprehensive water-wire connectivity *φ*_ow_ was quantified as the smooth geometric average of the mean occupancies between paired, adjacent virtual sites located across the pore lumen, as formulated in equation [3],

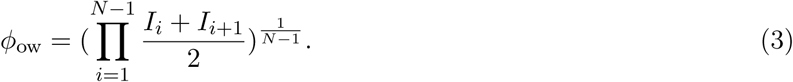

#### String Propagation

The strings are propagated from *i*-th iteration to (*i* + 1)-th iteration accroding to the following equation [4]:

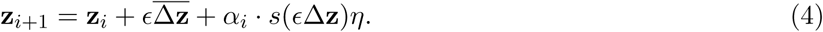

Here **z***_i_* and **z***_i_*_+1_ represent the image positions in *i*-th and *i* + 1-th iteration, respectively. The term 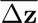 represents the mean image displacement Δ**z** within the CV space sampled during the swarm trajectories, while *ɛ* serves as a regularization parameter for numerical smoothing. The standard deviation of the image displacement Δ**z** within the CV space during swarm sampling is given by *s*(Δ**z**). *η* is defined as a stochastic variable drawn from a standard normal distribution, *η* ∼ *N*(0, 1), and *α_i_* = (1 − *i/i*_0_)*α* represents a linear annealing factor that decays to zero at the target iteration threshold *i*_0_ to facilitate sampling. This stochastic white-noise term introduces controlled thermal fluctuations, which assist string propagation in exploring a wider ensemble of configurations in the CV space toward the MFEP. For the simulations reported in this work, the parameters were employed as *ɛ* = 1.0, *α* = 5.0 and *i*_0_ = 25.

### First-order Electrodiffusion Approximation of Permeation Barrier from Conductance

#### Homogeneous Diffusion Model

We model ion permeation through the MERS ETM as a diffusion process governed by an intrinsic potential of mean force *F* (*z*) and an external electric potential Φ(*z*) along the pore axis *z* (oriented from the N-terminus to the C-terminus), the net ionic flux *J* is described by the Nernst-Planck equation:

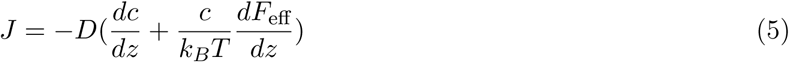

Here, *D* represents the average effective intra-pore diffusion coefficient, and *F*_eff_ = *F* (*z*) + *qe*Φ(*z*) is the effective potential experienced by an ion with valence *q*. The electrolyte concentrations at the cisand transsides of the membrane are denoted as *c*_0_ and *c_L_*, respectively. Assuming a steady-state flux, integrating Equation 5 over the pore length *L* yields:

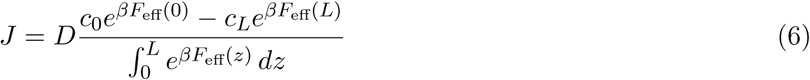

where *β* = 1*/k_B_T*. Setting the net flux *J* = 0 yields the equilibrium (reversal) potential, *V*_rev_ = 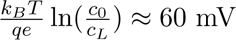. Experimental observations indicate that the pore conductance remains constant within the -30 mV to 150 mV range. For simplicity, we consider a membrane potential difference ΔΦ = Φ(*L*) −Φ(0) slightly above the equilibrium potential, defined as ΔΦ = *V*_rev_ + *δV*. Substituting this relation into Equation 6 gives:

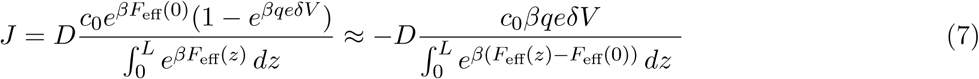

which holds in the limit of small 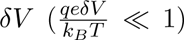. By introducing the dimensionless pore coordinate *ξ* = *z/L*, the ratio between intra-pore diffusion coefficient and its infinite-dilute limit *α* = *D/D_∞_*, the integral in the denominator can be scaled as 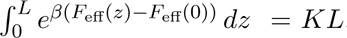, where 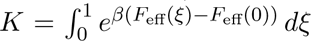. The pore conductance *γ* is consequently expressed as:

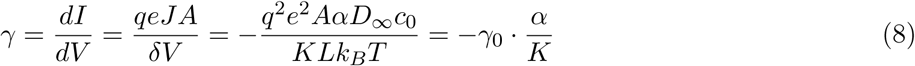

where 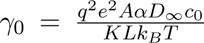 represents the conductance in the fully diffusion controlled regime. For the MERS ETM, the pore cross-sectional area is estimated as *A* = *πr*^2^ with average radius *r* ∼ 0.2 nm and an effective length *L* ∼ 4 nm. Under bulk concentrations of *c*_0_ = 500 mM and *c_L_* = 50 mM, substituting these parameters into the above equation of *γ*_0_ yields *γ*_0_ = 118 pS. This baseline value agrees remarkably well with the experimental measurement of *γ* = 113 ± 12.5 pS, implying that the dimensionless transport-partitioning factor *α/K* approaches unity. The desolvation penalties and other pore-related interactions captured by the partition coefficient *K* are comparable to, or only slightly exceed, the pore environment’s impact on the local diffusion coefficient captured by *α*. This demonstrates that ion permeation resides within a diffusioncontrolled regime regardless of the specific shape of the underlying PMF profile. The effective free-energy barrier is therefore bounded at only a few *k_B_T* which can be overcome by thermal fluctuation.

#### Inhomogeneous Diffusion Model

To account for spatial variations in transport dynamics, ion permeation can be modeled using a positiondependent diffusion coefficient, *D*(*z*). The intra-pore one-dimensional diffusion coefficient is estimated from local trajectory fluctuations along the pore axis via [1]

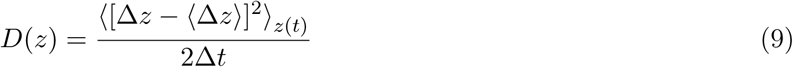

As shown in Fig. S5, the calculated *D*(*z*) for K^+^ and its ratio ( *α*) to the bulk diffusion coefficient at infinite dilution ranges from 0.1 to 1.7. Within this framework, the generalized Nernst–Planck equation takes the following form:

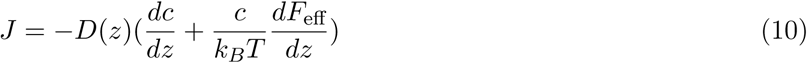

Assuming a steady-state flux and a membrane potential difference close to the equilibrium potential, an identical derivation applied to Eqs. 6–8 yields the single-channel conductance *γ*:

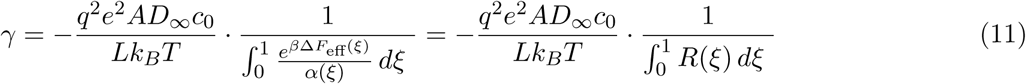

Here, *ξ* = *z/L* defines the dimensionless pore coordinate along the channel axis, and *α*(*ξ*) = *D*(*ξ*)*/D_∞_* represents the ratio of the local one-dimensional diffusion coefficient *D*(*z*) to the bulk diffusion coefficient at infinite dilution *D_∞_*. The integrand 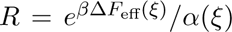 functions as a local resistivity; local drops in *D*(*z*) or elevations in the effective potential *F*_eff_ induce pronounced spikes in *R*, making it a robust metric for identifying gating effects. For simplicity, we assume an identical effective potential profile for both Na^+^ and K^+^, though in reality, the effective potential landscape for K^+^ is expected to be systematically lower and smoother. The corresponding profiles for *α*, *F*_eff_ and *R* are illustrated in Fig. S7. Notably, prominent spikes near residues F19, S30, and F33 indicate a substantial gating effect near these locations. Evaluating the spatial integral of *R* yields integrated resistivities of 2.0 × 10^4^ for Na^+^ and 1.4 × 10^4^ for K^+^. Substituting these integrated quantities into Equation 10 yields simulation predicted single-channel conductances of *γ* = 3.9 × 10*^−^*^3^ pS for Na^+^ and *γ* = 8.2 × 10*^−^*^3^ pS for K^+^. These are substantially lower than the experimental conductance, suggesting that the structural model and the underlying potential function require further improvements, which is not uncommon for a quantitative analysis of also the canonical ion channels[2–4].

### Benchmarking Cation-***π*** Interactions

#### Simulation Setup

To benchmark cation–*π* interactions, single-point energy calculations were performed on both representative snapshots from finite-temperature string (FTS) simulations and model systems. Conformations were sampled from image frames 20 to 26 of both additive CHARMM36m and polarizable Drude FTS trajectories. For these configurations, the coordinates of the phenylalanine (Phe33) side chain and the coordinating Na^+^ ion were extracted. To prevent valence artifacts, the truncated C*_β_* atom of the side chain was saturated with a hydrogen atom. Analogous model compounds were constructed by scanning a Na^+^ ion along geometric axes relative to the benzene ring center of a hydrogen-saturated Phe side chain.

Single-point energies were calculated using classical force fields (CHARMM36m and Drude) and a quantum mechanical (QM) method. Classical calculations were executed in OpenMM, employing self-consistent field (SCF) integration for the Drude model to ensure accurate polarization[5]. A hard upper limit of 0.02 nm on the displacement between the Drude particles and their parent atoms was applied to truncate SCF and prevent over-polarization artifacts. QM calculations were conducted via Density Functional Theory (DFT) in ORCA[6], utilizing the range-separated hybrid functional *ω*B97X-D[7] in conjunction with the def2-TZVP basis set[8]. Cation–*π* interaction energies (Δ*E*_int_) were evaluated as the difference between the total energy of the Na^+^-Phe complex and the sum of the isolated monomers. To eliminate basis set superposition error (BSSE) in the QM calculations, the Boys–Bernardi counterpoise correction[9] was applied using ghost atoms.

---

**Figure S1:**
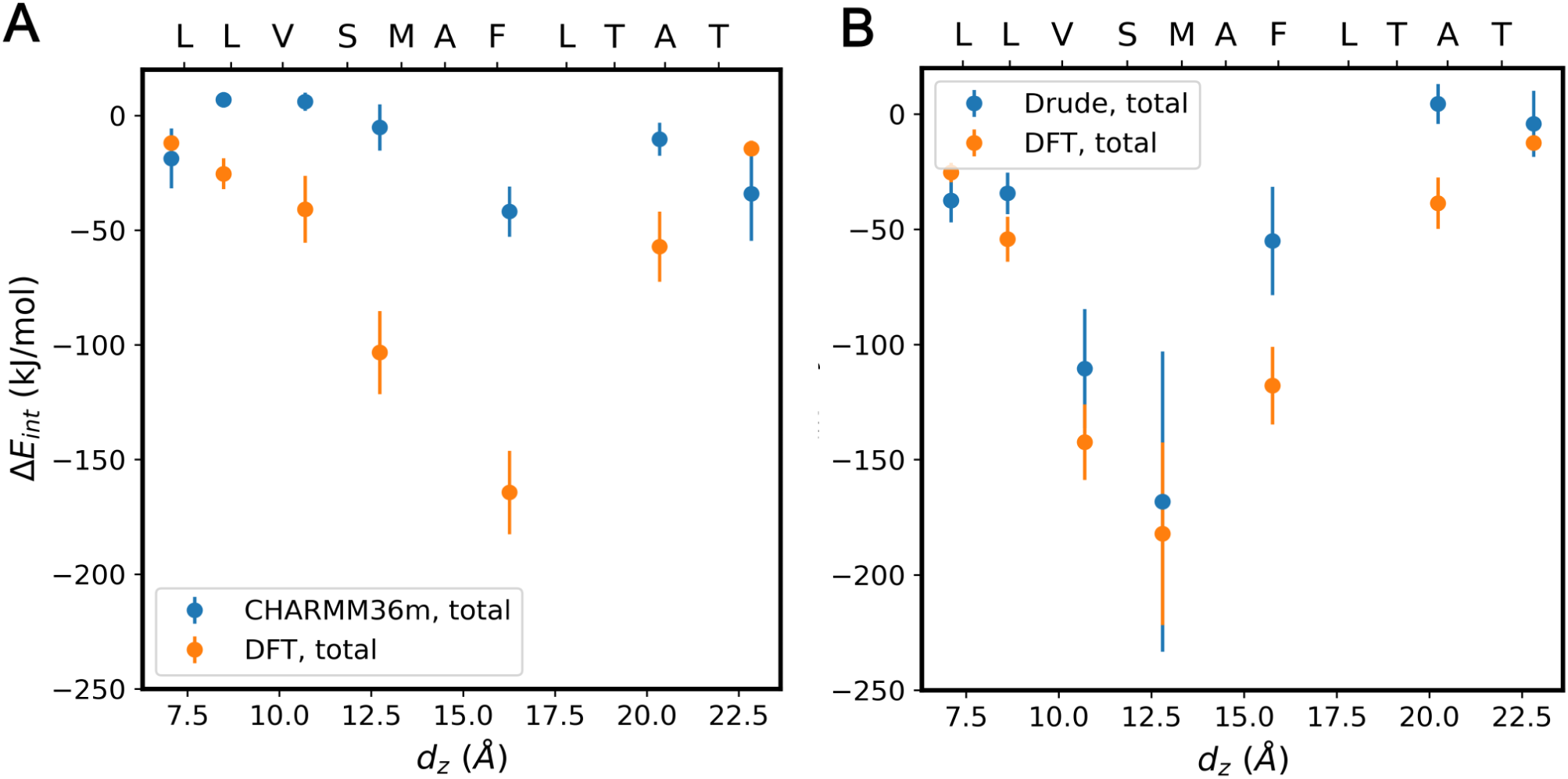
Comparison of cation–*π* interaction energies (Δ*E*_int_) calculated using classical force fields (A) fixed-charge CHARMM36m and (B) polarizable Drude and Density Functional Theory (DFT) at the *ω*B97X-D/def2-TZVP level. Configuration samples were extracted from finite-temperature string (FTS) simulation trajectories. All DFT calculations include Boys–Bernardi counterpoise corrections to account for basis set superposition error (BSSE). MERS ETM residue locations along the channel axis are indicated by their respective C*_α_* positions.

**Figure S2:**
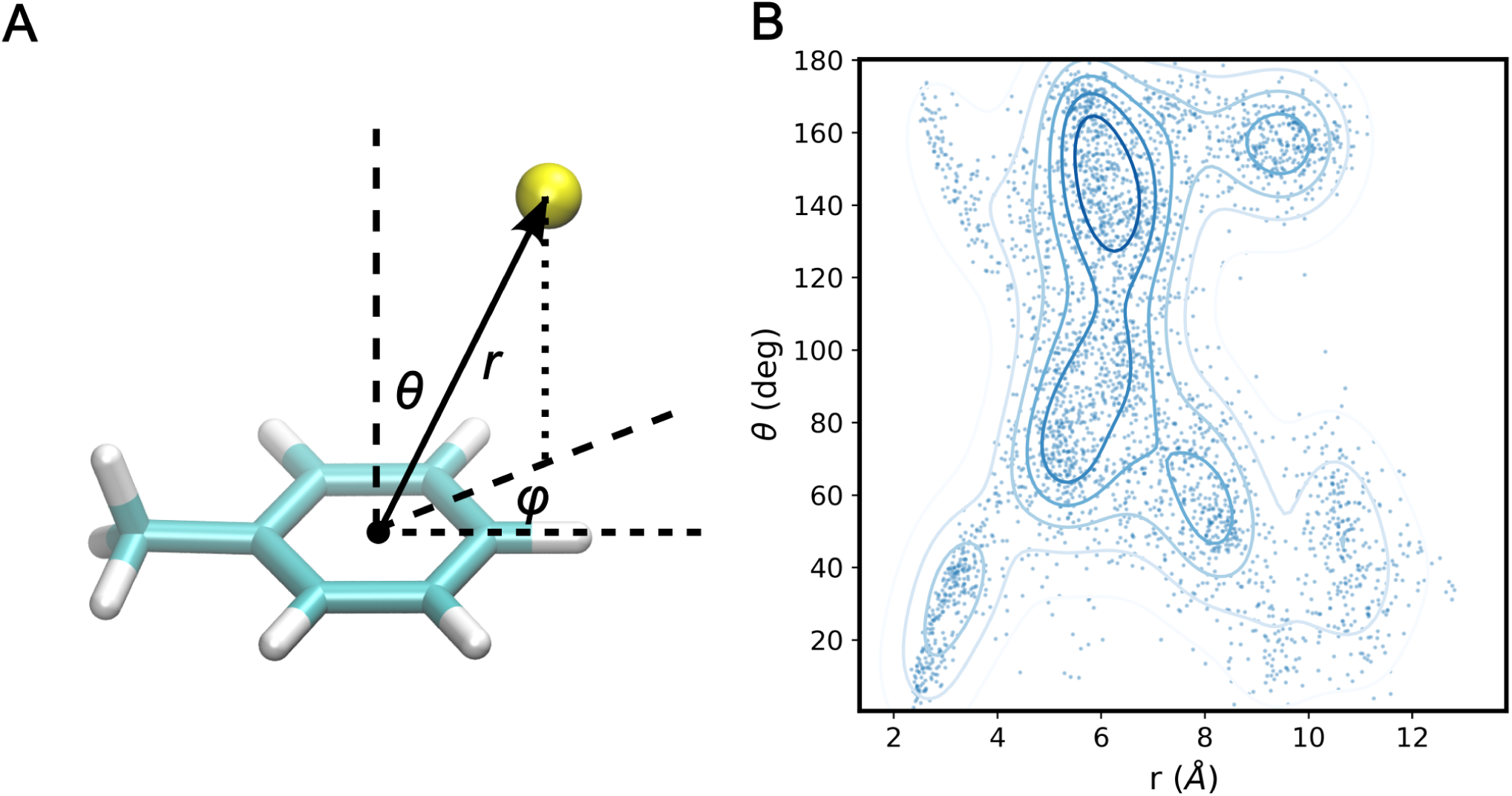
Structural coordinates and geometric distributions of the Na^+^–phenylalanine complex. (A) Schematic definition of the scanning coordinates and directions relative to the Phe side-chain benzene ring used to construct the model compound profiles. (B) Kernel density estimation (KDE) of the (r, *θ*) parameter space sampled from the selected Drude configurations during finite-temperature string (FTS) simulations, with probability density isosurfaces plotted up to the 90th percentile.

**Figure S3:**
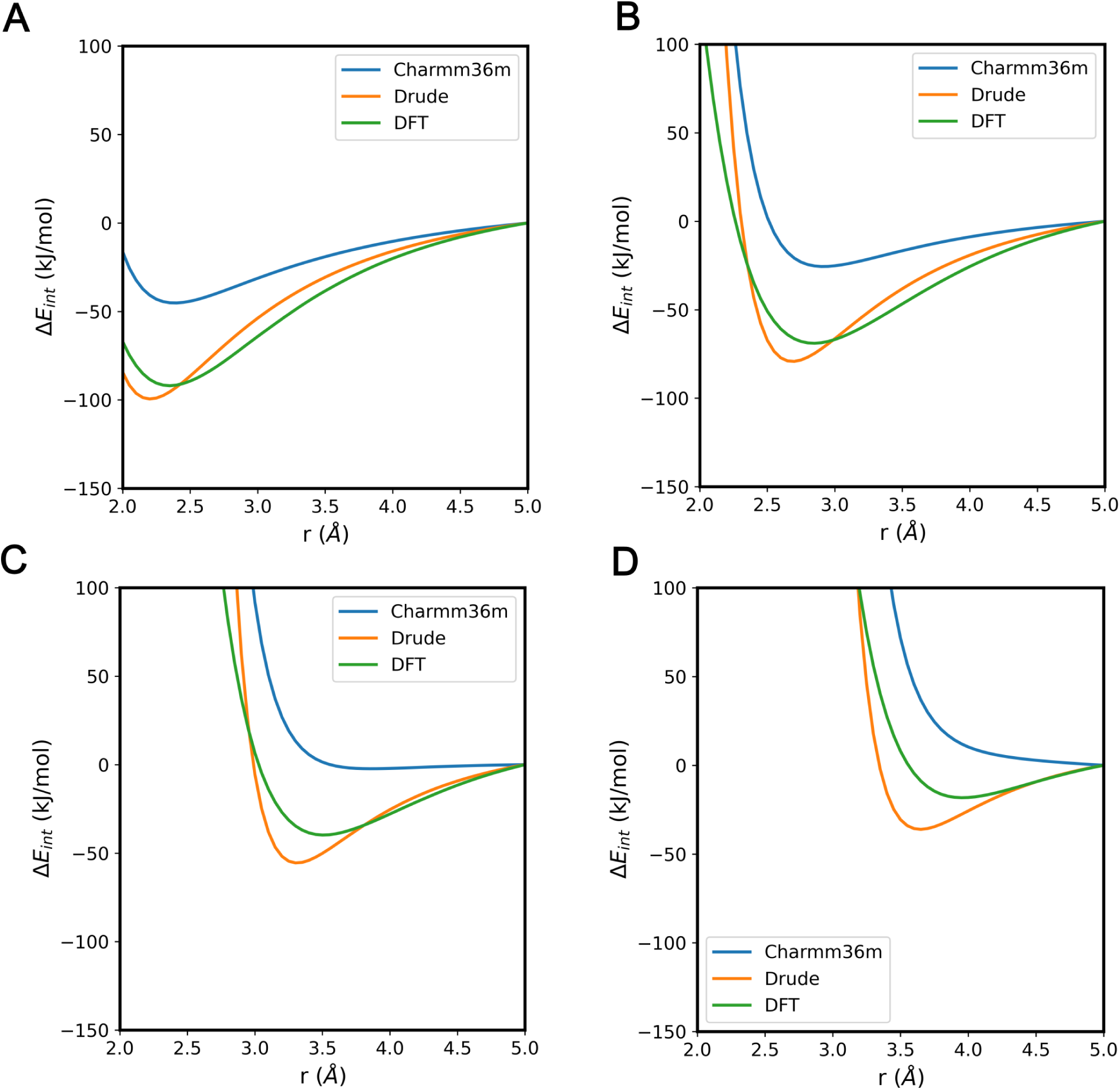
Cation–*π* interaction energies (Δ*E*_int_) calculated for the model Na^+^–phenylalanine complex across distinct spatial trajectories. Profiles are evaluated as a function of the Na^+^ scanning coordinate along varying angular configurations defined by the tilt (*θ*) and azimuthal (*φ*) angles relative to the benzene ring: (A) *θ* = 0*^◦^*; (B) *θ* = 30*^◦^, φ* = 30*^◦^*; (C) *θ* = 60*^◦^, φ* = 30*^◦^*; and (D) *θ* = 90*^◦^, φ* = 30*^◦^*.

**Figure S4:**
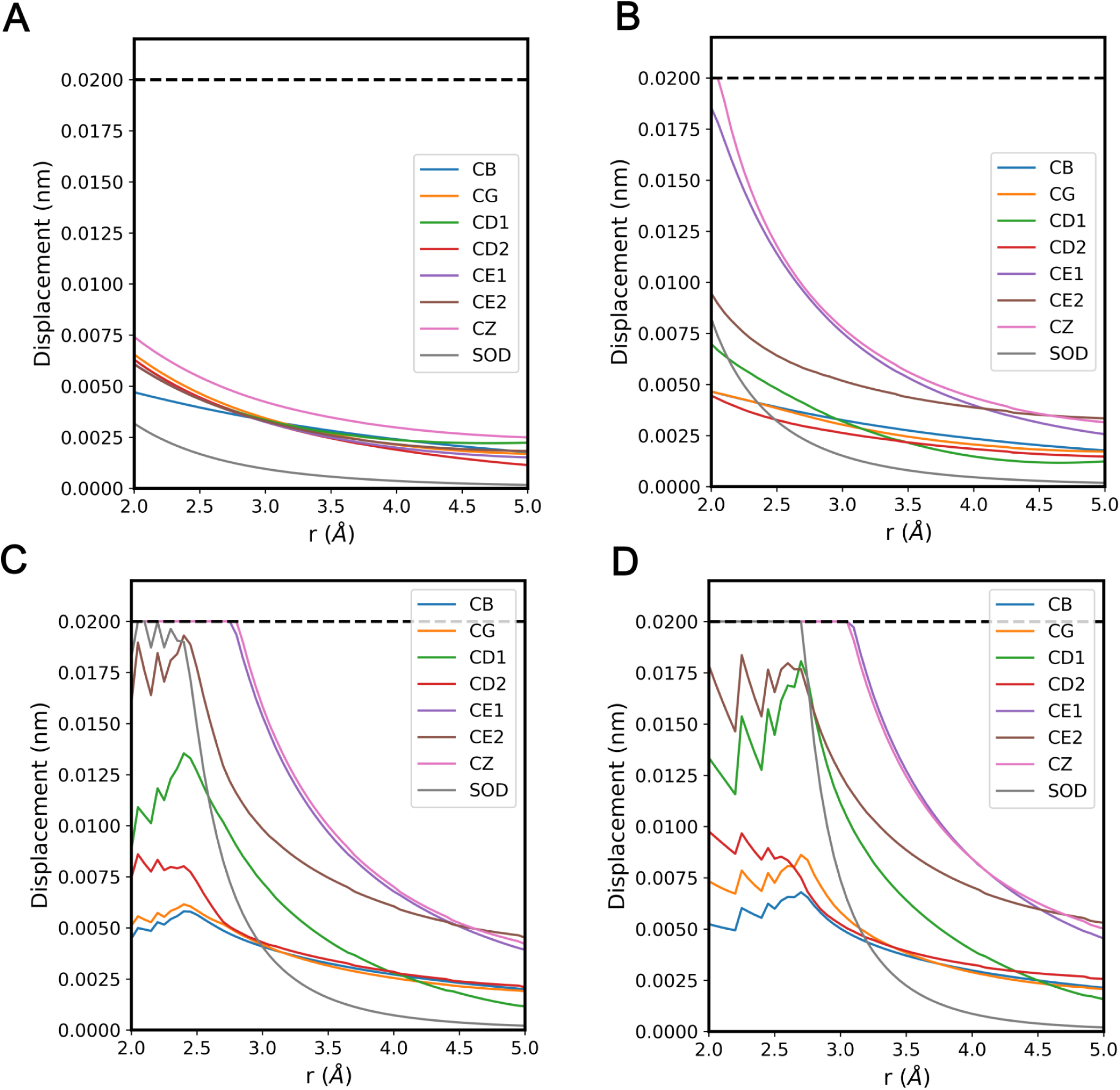
Displacement of Drude particles from their respective parent atoms calculated for the model Na^+^–phenylalanine complex across distinct spatial trajectories. Hard limit of 0.02 nm is highlighted in black dashed line. Profiles are evaluated as a function of the Na^+^ scanning coordinate along varying angular configurations defined by the tilt (*θ*) and azimuthal (*φ*) angles relative to the benzene ring: (A) *θ* = 0*^◦^*; (B) *θ* = 30*^◦^, φ* = 30*^◦^*; (C) *θ* = 60*^◦^, φ* = 30*^◦^*; and (D) *θ* = 90*^◦^, φ* = 30*^◦^*.

**Figure S5:**
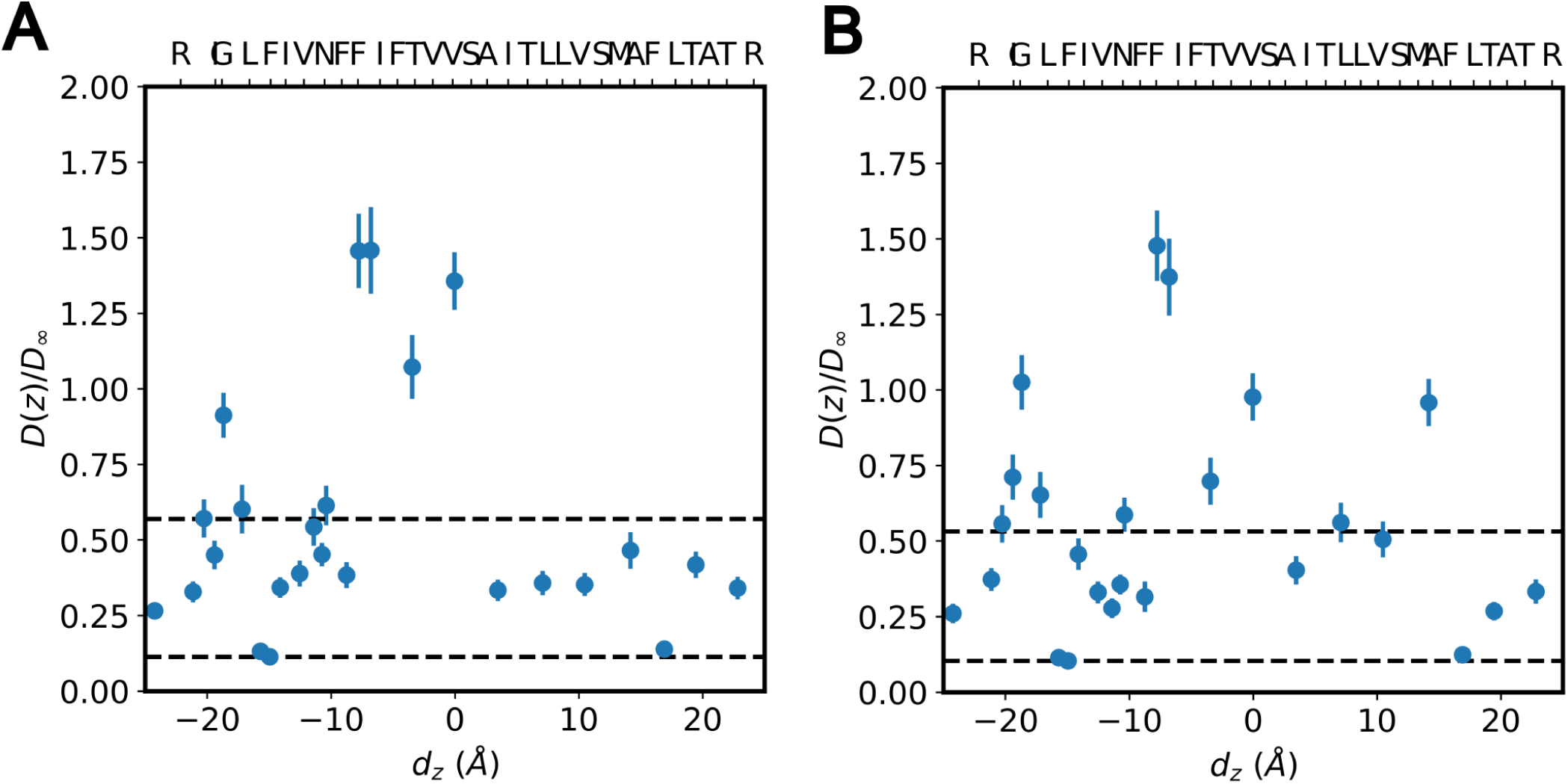
Local 1D diffusion coefficient *D*(*z*) along the pore axis. Normalized profiles are shown for (A) Na^+^ and (B) K^+^, where values are divided by the corresponding bulk diffusion coefficient at the infinite dilution limit *D_∞_* (Na^+^ : 1.33 × 10*^−^*^9^ m^2^ · s*^−^*^1^, K^+^ : 1.93 × 10*^−^*^9^ m^2^ · s*^−^*^1^). Data points are derived from 300 independent replicas at each image sampling Δ*t* = 1 ps, with standard deviations computed using a bootstrapping protocol (10,000 resamples with replacement). Black dashed lines highlight the minimum and average values for each profile (Na^+^: minimum 0.11, average 0.55; K^+^: minimum 0.10, average 0.53). MERS ETM residue locations along the channel axis are indicated by their respective C*_α_* positions.

**Figure S6:**
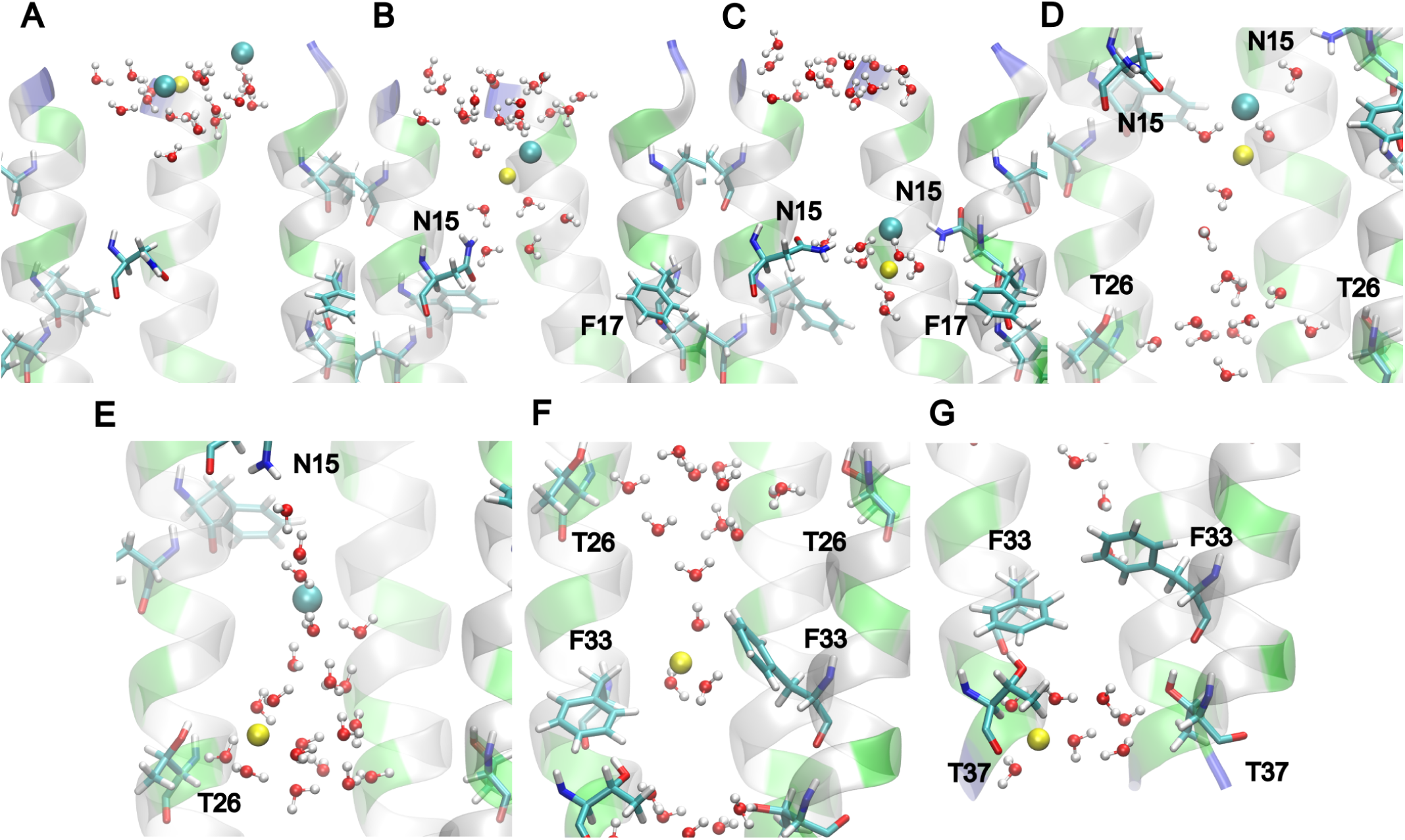
Representative configurations of Na^+^ permeation through the MERS ETM pore via the chaperoned ion-pair mechanism. Snapshots were obtained using the polarizable Drude force field. The permeating Na^+^ cation and its accompanying counterion Cl*^−^* are displayed as yellow and cyan spheres, respectively. Sequential panels depict key states along the pathway: (A) reactant state, (B) transition state 1 (TS_1_), (C) intermediate 1 (Int_1_), (D) transition state 2 (TS_2_), (E) intermediate 2 (Int_2_), (F) transition state 3 (TS_3_), and (G) product state. Critical channel residues, including aromatic residues (F12, F19, F33) and polar residues (N15, T26), are highlighted using the same color scheme as in Figure 1 (polar in green, hydrophobic in gray). Views are magnified to emphasize the local interactions between permeating ions, coordinating water molecules, and nearby protein residues.

**Figure S7:**
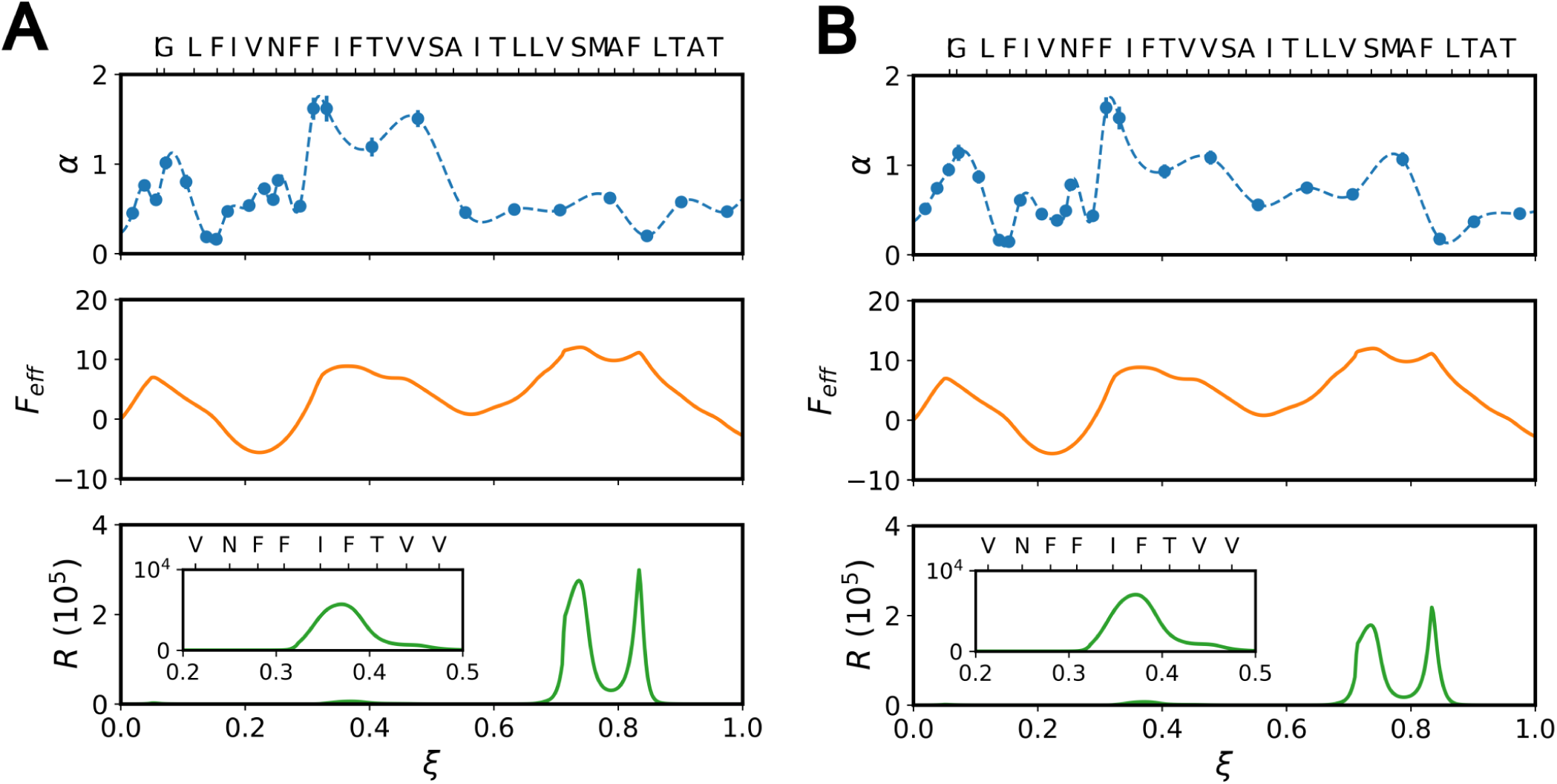
Diffusion and free-energy profiles along the pore axis. Local transport properties and potential of mean force (PMF) are plotted against the dimensionless pore coordinate *ξ* for (A) Na^+^ and (B) K^+^. Panels from top to bottom display: the ratio *α* of the local diffusion coefficient to the bulk diffusion coefficient at infinite dilution *D_∞_* (Na^+^ : 1.33 × 10*^−^*^9^ m^2^ · s*^−^*^1^; K^+^ : 1.93 × 10*^−^*^9^ m^2^ · s*^−^*^1^), where dashed lines denote the cubic spline interpolations utilized in subsequent local resistivity integrations; the effective potential *F*_eff_ in *k_B_T* (as mentioned in the text, here we apply the computed PMF for Na^+^ to both ions); and the local resistivity *R*. MERS ETM residue locations along the channel axis are indicated by their respective C*_α_* positions. (Insets) Detailed views of local resistivity *R* near the N-terminal phenylalanine cluster (F16, F17, and F19) highlighting the gating effect.

## References

[1] Miao Yu, Hans H. Funke, John L. Falconer, and Richard D. Noble. Gated ion transport through dense carbon nanotube membranes. Journal of the American Chemical Society, 132(24):8285–8290, May 2010.

[2] Hui Li, Joseph S. Francisco, and Xiao Cheng Zeng. Unraveling the mechanism of selective ion transport in hydrophobic subnanometer channels. Proceedings of the National Academy of Sciences, 112(35):10851– 10856, Aug 2015.

[3] J. W. Polster, F. Aydin, J. P. de Souza, M. Z. Bazant, T. A. Pham, and Z. S. Siwy. Rectified and salt concentration dependent wetting of hydrophobic nanopores. J. Am. Chem. Soc., 144:11693–11705, 2022.

[4] G. Hummer, J. C. Rasaiah, and J. P. Noworyta. Water conduction through the hydrophobic channel of a carbon nanotube. Nature, 414:188–190, 2001.

[5] J. Geng, K. Kim, J. F. Zhang, A. Escalada, R. Tunuguntla, L. R. Comolli, F. I. Allen, A. V. Shnyrova, K. R. Cho, D. Munoz, Y. M. Wang, C. P. Grigoropoulos, C. M. Ajo-Franlin, V. A. Frolov, and A. Noy. Stochastic transport through carbon nanotubes in lipid bilayers and live cell membranes. Nature, 514:612, 2014.

[6] Fangqiang Zhu and Gerhard Hummer. Drying transition in the hydrophobic gate of the glic channel blocks ion conduction. Biophysical Journal, 103(2):219–227, July 2012.

[7] C. I. Lynch, S. Rao, and M. S. P. Sansom. Water in nanopores and biological channels: A molecular simulation perspective. Chem. Rev., 120:10298–10335, 2020.

[8] D. Seiferth, P. C. Biggin, and S. J. Tucker. When is a hydrophobic gate not a hydrophobic gate? J. Gen. Physiol., 154:e202213210, 2022.

[9] Z. Jia, M. Yazdani, G. Zhang, J. Cui, and J. Chen. Hydrophobic gating in BK channels. Nat. Commun., 9:3408, 2018.

[10] Jiahua Deng and Qiang Cui. Electronic polarization leads to a drier dewetted state for hydrophobic gating in the big potassium channel. The Journal of Physical Chemistry Letters, 15(29):7436–7441, July 2024.

[11] R. H. Tunuguntla, F. I. Allen, K. Kim, A. Belliveau, and A. Noy. Ultrafast proton transport in sub-1-nm diameter carbon nanotube porins. Nat. Nanotechnol., 11:639, 2016.

[12] João Medeiros-Silva, Aurelio J. Dregni, Noah H. Somberg, Pu Duan, and Mei Hong. Atomic structure of the open sars-cov-2 e viroporin. Science Advances, 9(41), Oct 2023.

[13] Venkata S. Mandala, Matthew J. McKay, Alexander A. Shcherbakov, Aurelio J. Dregni, Antonios Kolocouris, and Mei Hong. Structure and drug binding of the sars-cov-2 envelope protein transmembrane domain in lipid bilayers. Nature Structural I&; Molecular Biology, 27(12):1202–1208, Nov 2020.

[14] Iva Sučec, Bingqing Xia, Noah H. Somberg, Yi Wang, Hyunil Jo, Shuangqu Li, Barbara Perrone, Zhaobing Gao, and Mei Hong. Ion channel structure and function of the mers coronavirus e protein. Science Advances, 11(28), July 2025.

[15] E. Gouaux and R. MacKinnon. Principles of selective ion transport in channels and pumps. Science, 310:1461–1465, 2005.

[16] B. Roux. Ion conduction and selectivity in k^+^ channels. Annu. Rev. Biophys. Biomol. Struct., 34:153– 171, 2005.

[17] João Medeiros-Silva, Yanina Pankratova, Iva Sučec, Aurelio J. Dregni, and Mei Hong. Polar networks mediate ion conduction of the sars-cov-2 envelope protein. Journal of the American Chemical Society, 147(1):746–757, Dec 2024.

[18] I. Sucec, D. Tymochko, Z. Wan, Q. Cui, and M. Hong. Aromatic residues regulate potassium-ion induced conformational selection and hydration of the mers coronavirus e viroporin. Submitted, 2026.

[19] Weinan E, Weiqing Ren, and Eric Vanden-Eijnden. Finite temperature string method for the study of rare events. The Journal of Physical Chemistry B, 109(14):6688–6693, Feb 2005.

[20] Eric Vanden-Eijnden and Maddalena Venturoli. Revisiting the finite temperature string method for the calculation of reaction tubes and free energies. The Journal of Chemical Physics, 130(19), May 2009.

[21] Ai Koizumi, Hirofumi Tahara, Tomonori Hirano, and Akihiro Morita. Revealing transient shuttling mechanism of catalytic ion transport through liquid–liquid interface. The Journal of Physical Chemistry Letters, 11(4):1584–1588, Feb 2020.

[22] Akihiro Morita, Ai Koizumi, and Tomonori Hirano. Recent progress in simulating microscopic ion transport mechanisms at liquid–liquid interfaces. The Journal of Chemical Physics, 154(8), Feb 2021.

[23] Kaito Naka, Teppei Kamimura, Tomonori Hirano, and Akihiro Morita. Free-energy analysis for facilitated ion transfer of multivalent ions at the liquid–liquid interface. The Journal of Physical Chemistry B, 129(23):5820–5830, June 2025.

[24] Chenghan Li and Gregory A. Voth. A quantitative paradigm for water-assisted proton transport through proteins and other confined spaces. Proceedings of the National Academy of Sciences, 118(49), Dec 2021.

[25] Benoît Roux, Toby Allen, Simon Bernèche, and Wonpil Im. Theoretical and computational models of biological ion channels. Quarterly Reviews of Biophysics, 37(1):15–103, Feb 2004.

[26] Hui Li, Janamejaya Chowdhary, Lei Huang, Xibing He, Alexander D. MacKerell, and Benoît Roux. Drude polarizable force field for molecular dynamics simulations of saturated and unsaturated zwitterionic lipids. Journal of Chemical Theory and Computation, 13(9):4535–4552, August 2017.

[27] Jay W. Ponder, Chuanjie Wu, Pengyu Ren, Vijay S. Pande, John D. Chodera, Michael J. Schnieders, Imran Haque, David L. Mobley, Daniel S. Lambrecht, Robert A. DiStasio, Martin Head-Gordon, Gary N. I. Clark, Margaret E. Johnson, and Teresa Head-Gordon. Current status of the amoeba polarizable force field. The Journal of Physical Chemistry B, 114(8):2549–2564, Feb 2010.

[28] Yue Shi, Zhen Xia, Jiajing Zhang, Robert Best, Chuanjie Wu, Jay W. Ponder, and Pengyu Ren. Polarizable atomic multipole-based amoeba force field for proteins. Journal of Chemical Theory and Computation, 9(9):4046–4063, Aug 2013.

[29] J. A. Lemkul, J. Huang, B. Roux, and A. D. MacKerell Jr. An empirical polarizable force field based on the classical Drude oscillator model: Development history and recent applications. Chem. Rev., 116:4983–5013, 2016.

[30] Yalun Yu, Richard M. Venable, Jonathan Thirman, Payal Chatterjee, Anmol Kumar, Richard W. Pastor, Benoît Roux, Alexander D. MacKerell, and Jeffery B. Klauda. Drude polarizable lipid force field with explicit treatment of long-range dispersion: Parametrization and validation for saturated and monounsaturated zwitterionic lipids. Journal of Chemical Theory and Computation, 19(9):2590–2605, Apr 2023.

[31] Jing Huang, Sarah Rauscher, Grzegorz Nawrocki, Ting Ran, Michael Feig, Bert L De Groot, Helmut Grubmüller, and Alexander D MacKerell Jr. Charmm36m: an improved force field for folded and intrinsically disordered proteins. Nature Methods, 14(1):71–73, 2017.

[32] M. Born. Volumen und hydratationswärme der ionen. Zeitschrift für Physik, 1(1):45–48, Feb 1920.

[33] Edward G. Hohenstein and C. David Sherrill. Wavefunction methods for noncovalent interactions. WIREs Computational Molecular Science, 2(2):304–326, July 2011.

[34] Joanna C. Flick, Dmytro Kosenkov, Edward G. Hohenstein, C. David Sherrill, and Lyudmila V. Slipchenko. Accurate prediction of noncovalent interaction energies with the effective fragment potential method: Comparison of energy components to symmetry-adapted perturbation theory for the s22 test set. Journal of Chemical Theory and Computation, 8(8):2835–2843, July 2012.

[35] Fangqiang Zhu and Gerhard Hummer. Pore opening and closing of a pentameric ligand-gated ion channel. Proceedings of the National Academy of Sciences, 107(46):19814–19819, Nov 2010.

[36] S. Chakrabarty and A. Warshel. Capturing the energetics of water insertion in biological systems: The water flooding approach. Proteins: Struct., Funct., & Bioinf., 81:93–106, 2013.

[37] P. Goyal, J. Lu, S. Yang, M. R. Gunner, and Q. Cui. Changing hydration level in an internal cavity modulates the proton affinity of a key glutamate in cytochrome c oxidase. Proc. Natl. Acad. Sci. U.S.A., 110:18886–18891, 2013.

[38] R. B. Liang, J. M. J. Swanson, Y. X. Peng, M. Wikström, and G. A. Voth. Multiscale simulations reveal key features of the proton-pumping mechanism in cytochrome c oxidase. Proc. Natl. Acad. Sci. U.S.A., 113:7420–7425, 2016.

[39] Chang Yun Son, Arun Yethiraj, and Qiang Cui. Cavity hydration dynamics in cytochrome c oxidase and functional implications. Proceedings of the National Academy of Sciences, 114(42), Oct 2017.

[40] Michael Röpke, Patricia Saura, Daniel Riepl, Maximilian C. Pöverlein, and Ville R. I. Kaila. Functional water wires catalyze long-range proton pumping in the mammalian respiratory complex i. Journal of the American Chemical Society, 142(52):21758–21766, Dec 2020.

[41] P. Chen, I. Vorobyov, B. Roux, and T. W. Allen. Molecular dynamics simulations based on polarizable models show that ion permeation interconverts between different mechanisms as a function of membrane thickness. J. Phys. Chem. B, 125:1020–1035, 2021.

[42] Gianni Klesse, Shanlin Rao, Stephen J. Tucker, and Mark S.P. Sansom. Induced polarization in molecular dynamics simulations of the 5-ht3 receptor channel. Journal of the American Chemical Society, 142(20):9415–9427, Apr 2020.

[43] Z. Jing, J. A. Rackers, L. R. Pratt, C. Liu, S. B. Rempe, and P. Ren. Thermodynamics of ion binding and occupancy in potassium channels. Chem. Sci., 12:8920–8930, 2021.

[44] C. Hui, R. de Vries, W. Kopec, and B. L. de Groot. Effective polarization in potassium channel simulations: Ion conductance, occupancy, voltage response, and selectivity. Proc. Natl. Acad. Sci. U.S.A., 122:e2423866122, 2025.

[45] Ramon Mendoza Uriarte and Benoît Roux. Conductance of a potassium channel in the limit of zero membrane potential. Biophysical Journal, 10.1016/j.bpj.2025.12.024, 2025.

[46] R. M. Uriarte and B. Roux. Force field sensitivity of ion occupancy in potassium channels. Biophys. J., 10.1016/j.bpj.2025.12.002, 2025.

[47] Sunhwan Jo, Taehoon Kim, Vidyashankara G. Iyer, and Wonpil Im. Charmm-gui: A web-based graphical user interface for charmm. Journal of Computational Chemistry, 29(11):1859–1865, June 2008.

[48] William L. Jorgensen, Jayaraman Chandrasekhar, Jeffry D. Madura, Roger W. Impey, and Michael L. Klein. Comparison of simple potential functions for simulating liquid water. The Journal of Chemical Physics, 79(2):926–935, July 1983.

[49] In Suk Joung and Thomas E. Cheatham. Determination of alkali and halide monovalent ion parameters for use in explicitly solvated biomolecular simulations. The Journal of Physical Chemistry B, 112(30):9020–9041, July 2008.

[50] Mark James Abraham, Teemu Murtola, Roland Schulz, Szilárd Páll, Jeremy C. Smith, Berk Hess, and Erik Lindahl. Gromacs: High performance molecular simulations through multi-level parallelism from laptops to supercomputers. SoftwareX, 1-2:19–25, 2015.

[51] GROMACS Development Team. Gromacs 2021 source code, Jan 2021.

[52] Giovanni Bussi, Davide Donadio, and Michele Parrinello. Canonical sampling through velocity rescaling. The Journal of Chemical Physics, 126(1):014101, 01 2007.

[53] Ulrich Essmann, Lalith Perera, Max L. Berkowitz, Tom Darden, Hsing Lee, and Lee G. Pedersen. A smooth particle mesh ewald method. The Journal of Chemical Physics, 103(19):8577–8593, Nov 1995.

[54] Berk Hess, Henk Bekker, Herman J. C. Berendsen, and Johannes G. E. M. Fraaije. Lincs: A linear constraint solver for molecular simulations. Journal of Computational Chemistry, 18(12):1463–1472, Sept 1997.

[55] Guillaume Lamoureux and Benoıt Roux. Modeling induced polarization with classical drude oscillators: Theory and molecular dynamics simulation algorithm. The Journal of Chemical Physics, 119(6):3025–3039, Aug 2003.

[56] Wei Jiang, David J. Hardy, James C. Phillips, Alexander D. MacKerell, Klaus Schulten, and Benoît Roux. High-performance scalable molecular dynamics simulations of a polarizable force field based on classical drude oscillators in namd. The Journal of Physical Chemistry Letters, 2(2):87–92, Dec 2010.

[57] Jing Huang, Justin A. Lemkul, Peter K. Eastman, and Alexander D. MacKerell. Molecular dynamics simulations using the drude polarizable force field on gpus with openmm: Implementation, validation, and benchmarks. Journal of Computational Chemistry, 39(21):1682–1689, May 2018.

[58] Peter Eastman, Raimondas Galvelis, Raúl P. Peláez, Charlles R. A. Abreu, Stephen E. Farr, Emilio Gallicchio, Anton Gorenko, Michael M. Henry, Frank Hu, Jing Huang, Andreas Krämer, Julien Michel, Joshua A. Mitchell, Vijay S. Pande, João PGLM Rodrigues, Jaime Rodriguez-Guerra, Andrew C. Simmonett, Sukrit Singh, Jason Swails, Philip Turner, Yuanqing Wang, Ivy Zhang, John D. Chodera, Gianni De Fabritiis, and Thomas E. Markland. Openmm 8: Molecular dynamics simulation with machine learning potentials. The Journal of Physical Chemistry B, 128(1):109–116, Dec 2023.

[59] The PLUMED consortium. Promoting transparency and reproducibility in enhanced molecular simulations. Nature Methods, 16(8):670–673, July 2019.

[60] Albert C. Pan, Deniz Sezer, and Benoît Roux. Finding transition pathways using the string method with swarms of trajectories. The Journal of Physical Chemistry B, 112(11):3432–3440, Feb 2008.

[61] B. Roux. String method with swarms-of-trajectories, mean drifts, lag time, and committor. J. Phys. Chem. A, 125:7558–7571, 2021.

[62] Paolo Raiteri, Alessandro Laio, Francesco Luigi Gervasio, Cristian Micheletti, and Michele Parrinello. Efficient reconstruction of complex free energy landscapes by multiple walkers metadynamics. The Journal of Physical Chemistry B, 110(8):3533–3539, Oct 2005.

[63] Alessandro Barducci, Giovanni Bussi, and Michele Parrinello. Well-tempered metadynamics: A smoothly converging and tunable free-energy method. Physical Review Letters, 100(2), Jan 2008.

[64] Davide Branduardi, Francesco Luigi Gervasio, and Michele Parrinello. From a to b in free energy space. The Journal of Chemical Physics, 126(5), Feb 2007.

[65] Grisell Díaz Leines and Bernd Ensing. Path finding on high-dimensional free energy landscapes. Physical Review Letters, 109(2), July 2012.

[66] Davide Branduardi, Giovanni Bussi, and Michele Parrinello. Metadynamics with adaptive gaussians. Journal of Chemical Theory and Computation, 8(7):2247–2254, June 2012.

## References

[1] Benoît Roux, Toby Allen, Simon Bernèche, and Wonpil Im. Theoretical and computational models of biological ion channels. Quarterly Reviews of Biophysics, 37(1):15–103, Feb 2004.

[2] Ramon Mendoza Uriarte and Benoît Roux. Conductance of a potassium channel in the limit of zero membrane potential. Biophysical Journal, 10.1016/j.bpj.2025.12.024, 2025.

[3] R. M. Uriarte and B. Roux. Force field sensitivity of ion occupancy in potassium channels. Biophys. J., 10.1016/j.bpj.2025.12.002, 2025.

[4] C. Hui, R. de Vries, W. Kopec, and B. L. de Groot. Effective polarization in potassium channel simulations: Ion conductance, occupancy, voltage response, and selectivity. Proc. Natl. Acad. Sci. U.S.A., 122:e2423866122, 2025.

[5] Jing Huang, Justin A. Lemkul, Peter K. Eastman, and Alexander D. MacKerell. Molecular dynamics simulations using the drude polarizable force field on gpus with openmm: Implementation, validation, and benchmarks. Journal of Computational Chemistry, 39(21):1682–1689, May 2018.

[6] Frank Neese. Software update: The orca program system—version 6.0. WIREs Computational Molecular Science, 15(2), Mar 2025.

[7] Jeng-Da Chai and Martin Head-Gordon. Long-range corrected hybrid density functionals with damped atom–atom dispersion corrections. Physical Chemistry Chemical Physics, 10(44):6615, 2008.

[8] Florian Weigend and Reinhart Ahlrichs. Balanced basis sets of split valence, triple zeta valence and quadruple zeta valence quality for h to rn: Design and assessment of accuracy. Physical Chemistry Chemical Physics, 7(18):3297, 2005.

[9] S.F. Boys and F. Bernardi. The calculation of small molecular interactions by the differences of separate total energies. some procedures with reduced errors. Molecular Physics, 19(4):553–566, Oct 1970.

